# OmniSyn unifies target-aware molecular generation and optimization within a synthesis-native LLM framework across the human proteome

**DOI:** 10.64898/2026.09.02.748775

**Authors:** Zheng Qin, Yuzhang Li, Yueqing Zhang, Yunshuo Zhao, Huan Yee Koh, Zheng Wan, Haocheng Ren, Changying Huang, Yiming Shi, Zhenguo Wu, Yaosen Min, Jing Yang, Xiao He, Duanhua Cao

## Abstract

Designing target-specific bioactive molecules with actionable synthesis routes for the human proteome holds enormous potential for expanding therapeutic discovery, but remains a challenge. Existing target-aware generative models often depend on protein structures and generate molecules before assessing synthetic feasibility.

Here we present OmniSyn, a protein-sequence-conditioned Mixture-of-Experts (MoE) language model that couples a task-conditioned interaction module with a synthesis-action decoder to generate molecules with explicit synthesis traces, thereby unifying de novo ligand generation, synthesizability projection and hit-to-lead (H2L) optimization within synthesis-traceable chemical space. OmniSyn is pre-trained with self-distillation and post-trained with task-specific reinforcement learning (RL) to adapt expert routing and optimize molecular properties across design modes. On unseen protein targets from MolGenBench, a real-world drug-discovery benchmark, OmniSyn achieves state-of-the-art performance across de novo design and H2L optimization, including target-awareness and hit-rediscovery metrics, despite relying only on protein sequences rather than three-dimensional (3D) pocket structures.

By embedding synthesis planning into the design process, OmniSyn shifts molecular generation from a generate-then-filter paradigm toward design-with-synthesis paradigm, transforming virtual predictions into experimentally actionable candidates. Independent AiZynthFinder evaluation yielded retrosynthetic success rates of 68.47% for de novo generation and 71.92% for H2L optimization, improving over the strongest baselines by 61.3% and 184.2%, respectively, and supporting the synthetic feasibility of OmniSyn-generated molecules. In synthesizability projection, OmniSyn further converts outputs from external generative models into close analogues with improved retrosynthetic feasibility while preserving molecular similarity. Having established strong benchmark performance and external retrosynthetic feasibility, we next applied OmniSyn at human-proteome scale, spanning more than 21,000 targets. Rapid sequence-conditioned sampling enabled the construction of, to our knowledge, the largest human-proteome-scale generative virtual library, comprising 2.7 billion target-specific molecules, each accompanied by model-derived synthesis traces and target-specific prioritization scores. By enabling scalable target-specific molecular design with synthesis-aware generation across the human proteome, OmniSyn opens new opportunities for exploring previously inaccessible therapeutic targets, including those lacking experimentally resolved structures.

## Introduction

Advances in computer vision, natural language processing and multimodal generation have expanded artificial intelligence from perception to reasoning and large-scale content creation^1,2^. These general advances have also accelerated the emergence of AI for science, where data-driven models are increasingly used not only to analyze observations, but also to predict, simulate and design complex systems^3^. In drug discovery, AI is increasingly used to identify targets, model perturbations and generate therapeutic candidates, yet molecular design remains limited by the need to produce compounds that are simultaneously target-relevant, chemically valid and synthetically accessible^4^. Recent advances have reshaped this landscape: AlphaFold and related methods have broadened access to protein structures^5,6^, protein generative models such as RFdiffusion^7^ and all-atom design frameworks have enabled de novo biomolecular design^8–10^ and shown that functional biomolecules can be designed directly rather than selected from pre-existing libraries. In parallel, large protein language models such as the ESM series^11,12^ have demonstrated that evolutionary and functional constraints can be learned from sequence alone, enabling scalable representations of proteins even when experimentally resolved structures are unavailable. More broadly, flow matching has emerged as a continuous-time generative paradigm, with applications across small molecules, proteins, biomolecular interactions and cellular states^13^. These advances are shifting drug discovery toward generative exploration of biological and chemical space. However, scalable target-conditioned molecular design with practical synthesis routes remains an unresolved challenge.

Target-aware small-molecule design remains central to drug discovery^4^. Recent structure-based drug design (SBDD) models such as Pocket2Mol^14^, DiffSBDD^15^, and PocketFlow^16^ have improved de novo ligand generation by conditioning on 3D pockets, but their reliance on accurate structures limits applicability to many disease-relevant target^17,18^. and their outputs often lack executable synthesis routes. As a result, generated molecules can be chemically attractive in silico but difficult to make, limiting their value for experimental validation. Sequence-based models such as ProtoBind-Diff^19^ and LaMGen^20^ reduce structural dependence, but have not yet consistently matched structure-based methods in target-aware accuracy^21^. Thus, scalable target-aware generation that combines protein-sequence conditioning with explicit synthesis planning remains an open challenge. A complementary line of work embeds synthetic feasibility directly into molecular generation^22^. Autoregressive approaches such as PrexSyn^23^ and SynFormer^24^ formulate molecular generation as sequential synthesis planning, in which molecules are represented through building-block and reaction decisions, whereas SynFlowNet^25^ explores reaction-valid trajectories using reward-directed sampling. ClickGen^26^ extends reaction-based generation to target-directed design, although its accessible chemical space is limited by a narrow set of transformations, primarily click reactions and amidation. By constructing molecules through predefined reactions, these approaches provide explicit synthesis traces rather than assessing synthetic feasibility only after generation. However, most existing reaction-based models are conditioned primarily on ligand representations, molecular properties, docking scores or task-specific oracles, rather than providing a general formulation for protein-conditioned generation^23–26^. Their scope is also commonly restricted to analogue generation, unconditional exploration, property optimization or narrow reaction vocabularies^27^. Consequently, reaction-based generation improves synthetic tractability but remains insufficiently integrated with scalable target-specific de novo design and H2L optimization.

H2L optimization is a central stage of small-molecule drug discovery. Rather than exploring chemical space without constraints, medicinal chemist typically refine known hits, privileged scaffolds or synthetically tractable fragments through iterative structure-activity relationship analysis and analogue synthesis to improve potency, selectivity and developability^28,29^. Generative models for this setting must therefore preserve activity-relevant molecular context while exploring target-specific substitutions and scaffold elaborations. Despite its practical importance, H2L optimization remains less developed than de novo generation in AI-driven molecular design^30^. These tasks are generally modelled separately, although both require target relevance, chemical validity and appropriate molecular properties, differing primarily in whether generation is additionally constrained by existing fragments. Models such as Delete^30^ and DiffDec^31^ support pocket-conditioned optimization through fragment manipulation or scaffold decoration, but remain dependent on 3D protein structures and separate from de novo, reaction-based generation. A shared formulation could expand the available supervision and facilitate transfer between discovery and optimization, consistent with recent evidence that de novo and fragment-constrained generation can benefit from multi-task training^32^. This gap is particularly evident for sequence-conditioned models, for which unified approaches spanning target-aware de novo design and synthesis-traceable H2L optimization remain scarce.

A complementary challenge arises when a molecule of interest is biologically promising but difficult to access synthetically, as may occur for structurally complex natural products or candidates proposed by generative models without reaction constraints. Synthesizability projection addresses this setting by seeking a structurally similar and synthetically tractable analogue that preserves the core features of the input molecule. Yet this capability is rarely integrated with target-aware de novo generation or H2L optimization. Taken together, these limitations expose a fragmented landscape. Structure-based models provide strong target conditioning but depend on accurate protein structures and commonly treat synthesizability as a downstream consideration. Reaction-based models generate molecules through valid transformations but generally lack broad target conditioning, whereas lead-optimization models preserve molecular context but remain task-specific and often require 3D pockets. This separation limits knowledge transfer across related design regimes. A unified model spanning target-aware de novo generation, synthesizability projection and H2L optimization would therefore address a central gap in molecular design.

Here we introduce OmniSyn, a protein-sequence-conditioned MoE language model trained through pre-training and post-training that unifies target-aware molecular generation, optimization and synthesis planning. OmniSyn couples a reaction-based autoregressive decoder^23^ with a novel Protein-Ligand Interaction Mixture-of-Experts (PLIM) module informed by pretrained protein^33^ and molecular models^34^. The decoder assembles molecules from more than 220,000 purchasable building blocks using 115 common reaction templates, providing an explicit synthesis trace for each completed molecule. PLIM integrates protein sequence representations, molecular context and task identity through token-level and pairwise representations, while task-conditioned expert routing adapts model computation across the three design tasks. OmniSyn is trained by self-distillation on high-quality pseudo synthesis trajectories, followed by reinforcement learning with task-specific objectives. This architecture unifies protein-sequence-conditioned de novo design, molecule-conditioned synthesizability projection and protein- and scaffold-conditioned H2L optimization within a reaction-valid generative framework.

We evaluated OmniSyn on MolGenBench^35^ across de novo design and H2L optimization. Despite conditioning target-aware generation on protein sequences rather than 3D pockets, OmniSyn outperformed nearly all structure-based baselines and achieved state-of-the-art target-awareness and hit-rediscovery performance. It attained scaffold and SMILES hit fractions of 2.5% and 0.255% in de novo design and corresponding hit rates of 10% and 0.163% in H2L optimization, with only 26% of targets in the lowest SMILES-level target-awareness range. OmniSyn also achieved the strongest synthetic-accessibility performance, including AiZynthFinder^36^ success rates of up to 73%, while providing explicit synthesis traces. In a separate projection task, it improved the synthetic feasibility of externally generated molecules while preserving structural similarity. Analyses of expert routing, protein-molecule pair representations and sparse-autoencoder features further revealed task specialization, localized interaction signals and chemically enriched decoder representations.

While conventional virtual screening is limited to ranking compounds within a static library, OmniSyn actively generates new target-conditioned molecules, thereby unlocking a significantly broader, synthesizable chemical space. DrugCLIP^37^ extended structure-based screening to approximately 10,000 human proteins and hundreds of millions of compounds, but remains constrained by a predefined library and requires predicted protein structures and pocket definitions. OmniSyn instead conditions directly on protein sequences, removing both requirements. We applied OmniSyn to more than 21,000 human proteins and constructed, to our knowledge, the largest target-specific generative virtual library, comprising 2.7 billion molecules. Each candidate is linked to an individual protein target and accompanied by a model-derived reaction route and target-specific prioritization scores, providing a searchable resource for proteome-wide target exploration and the prioritization of synthesis-ready candidates for experimental follow-up.

Together, OmniSyn unifies target-aware molecular generation and optimization within a sequence-conditioned, synthesis-native framework that does not require 3D pocket structures. Its extension across the human proteome yielded a 2.7-billion-molecule synthesis-traceable virtual library for target exploration and hit prioritization. More broadly, OmniSyn illustrates how molecular language models could evolve into unified design engines for closed-loop therapeutic discovery, jointly integrating target information, molecular transformation and synthesis to uncover new therapeutic molecules and target-ligand relationships, thereby expanding the opportunity to develop new treatment strategies for previously underexplored targets across the human proteome.

## Results

### Data construction and method overview

OmniSyn was developed as a unified sequence-conditioned model for reaction-based small-molecule design (**Fig. 1**). The final task tables were assembled from Papyrus^38^- and ChEMBL^39,40^-derived bioactivity records using task-specific, rather than uniform, selection rules^37–39^. Protein-conditioned de novo generation used target-ligand pairs linked to reviewed UniProt^41^ sequences. Molecule-only synthesizability projection used a canonical-SMILES deduplicated molecule set and supplied no protein information. H2L examples combined assay-derived chemical series with additional target-specific series; each example paired a protein sequence and an extracted maximum common substructure with related active molecules. Compounds were standardized and canonicalized with RDKit^42^, and the 35 MolGenBench unseen target identifiers used for evaluation were excluded from target-conditioned training tables to maintain consistency with the baseline models . The resulting task-specific data distributions are summarized in **Supplementary Figs. 1-3**, and complete filtering and dataset-construction details are provided in the **Supplementary Information**.

**Fig. 1.**
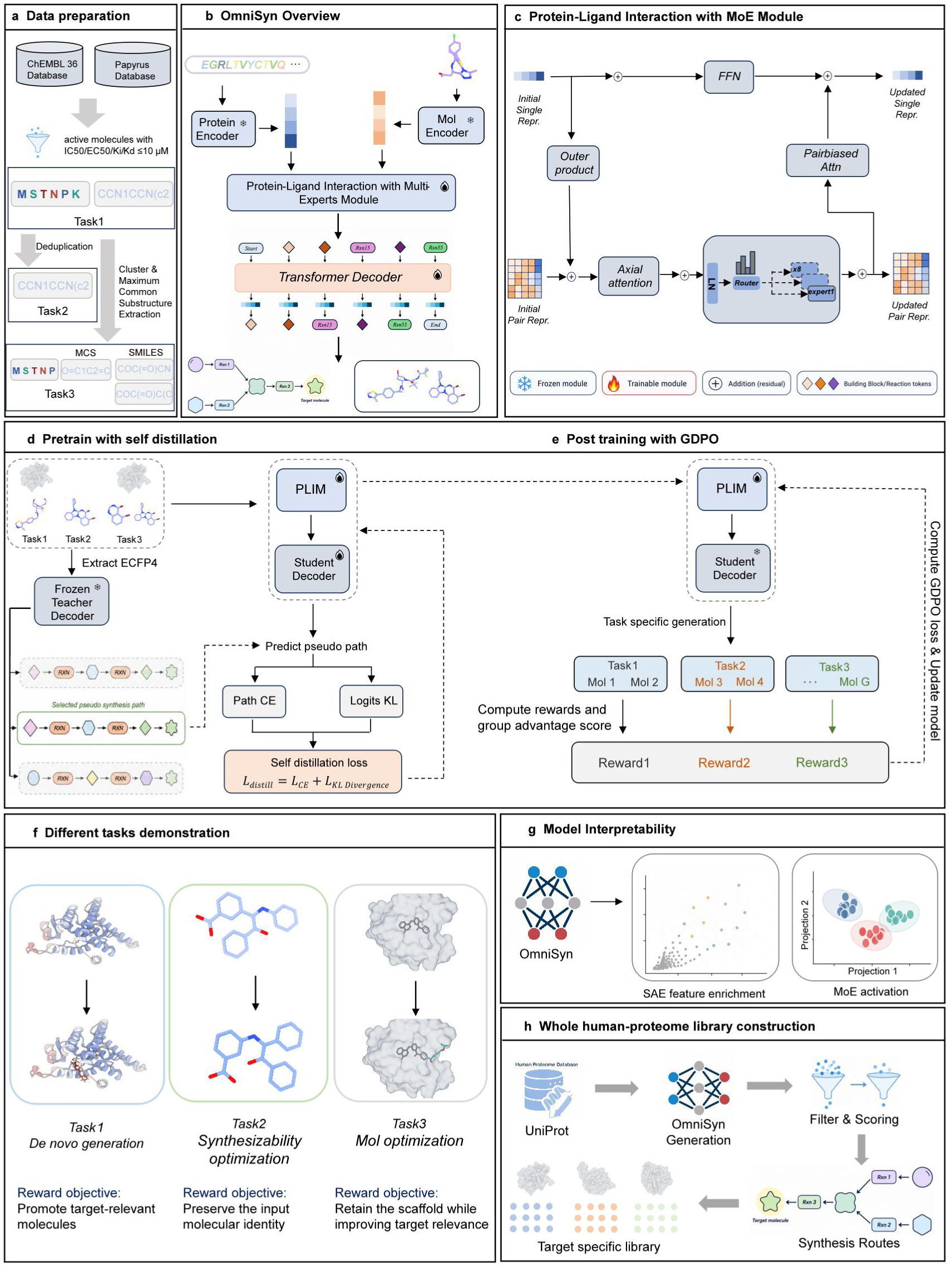
Overview of OmniSyn, its training strategy and downstream applications. **a**, Task-specific data construction and conditioning schemes for the three molecular-design settings. Papyrus and ChEMBL 36 records were processed using task-specific selection criteria, and target-associated records were mapped to reviewed UniProt sequences. The resulting examples were used for protein-conditioned de novo generation, molecule-only synthesizability projection and protein- and scaffold-conditioned H2L optimization. **b,** OmniSyn architecture, comprising frozen protein and molecular encoders, PLIM and the reaction-based OmniSyn decoder. **c,** A PLIM block. Pair initialization uses concatenated token pairs; the block itself contains a low-rank outer-product update, axial pair attention, top-two sparse expert routing, pair-biased attention and a single-token feed-forward update. **d,** Route self-distillation. A frozen base teacher supplies selected pseudo-paths; PLIM and decoder layers 0, 10 and 11 receive path cross-entropy and stop-gradient logit-KL supervision. **e,** GDPO post-training. Task-specific rewards are normalized separately and again after aggregation; PLIM is updated while the decoder is frozen. **f,** Input conditions and reward objectives for the three design tasks. **g,** Sparse-autoencoder and expert-routing analyses. **h,** Human-proteome library construction through sequence-conditioned generation, scoring and filtering, with retained entries linked to generated reaction routes. Flame and snowflake symbols denote trainable and frozen modules, respectively. SAE, sparse autoencoder; MCS, maximum common substructure; GDPO, Group Reward-Decoupled Normalization Policy Optimization.

OmniSyn integrates a decoder^23^ with a Protein-Ligand Interaction Mixture-of-Experts (PLIM) module (**Fig. 1b**). PLIM supplies task-specific conditioning representations. Protein sequences are encoded by frozen ESM-2^11^, whereas seed molecules or scaffolds are represented at atom level by a frozen molecular encoder^33^. Conditioned on these representations, the OmniSyn decoder autoregressively generates synthesis actions, including building-block selections and reaction-template decisions, that jointly specify the final product molecule and its explicit synthesis route. PLIM uses coupled single-token and pair representations to reason over the available task, protein and molecular context (**Fig. 1c**). A task token representing the task type and projected residue or atom embeddings form the single representation. Each ordered pair is initialized by concatenating its two single-token embeddings and applying a learnable linear projection. Within each PLIM block, a low-rank outer-product update transfers single-token information to the pair track; row-wise and column-wise axial attention then refine pair context^40^. A sparse MoE layer updates every valid pair position by task-biased top-two routing^41^. Pair-biased attention returns pair information to the single track. After four PLIM blocks, masked mean pooling and a two-layer conditioning projection produces a 1,024-dimensional condition vector used to autoregressively generates synthesis actions.

Training proceeded in two stages because a direct connection between a multimodal conditioner and the synthetic route decoder provides neither token-level alignment to the decoder’s established conditioning space nor an efficient signal for learning sparse task-dependent routing. In route self-distillation (**Fig. 1d**), a frozen route decoder teacher first generated candidate synthesis trajectories under ECFP4 conditioning; the best valid trajectory with ECFP4 similarity of at least 0.6 to the training molecule was cached as a pseudo-path. Teacher forcing these paths supplied dense action supervision before sequence-level reinforcement learning. For training efficiency, PLIM and only decoder layers 0, 10 and 11 were trainable during this stage. The remaining decoder parameters stayed fixed, preserving the reaction prior while adapting the condition interface. The self-distillation objective comprised two terms. A cross-entropy loss supervised each decoding step using the next action in the cached pseudo-path. A temperature-scaled KL divergence aligned the PLIM-conditioned action distribution with the stop-gradient distribution generated by the same decoder under ECFP4 conditioning. The self-distilled model was subsequently optimized with task-specific reinforcement learning (RL) (**Fig. 1e-f**). At this stage,the adapted OmniSyn decoder was frozen and only PLIM was updated, restricting policy improvement to conditional control rather than relearning reaction syntax. During the RL stage, we used Group Reward-Decoupled Normalization Policy Optimization (GDPO)^42^ to standardize each reward component within its rollout group before task-weighted aggregation, followed by a second normalization of the combined advantage. This procedure mitigates scale imbalance among heterogeneous rewards and preserves contributions from individual reward during multi-objective RL. The final objective combined clipped policy optimization, entropy regularization, a sampled-trajectory approximation to the reference-policy KL and the PLIM expert load-balancing term.

At inference, the task is selected by changing the input condition with a single set of model parameters. A protein sequence specifies de novo ligand generation; a molecule-only input specifies synthesizability projection; and a protein sequence together with a scaffold or fragment specifies H2L optimization. Each completed molecule is returned with its decoded reaction route. The trained model was subsequently examined through sparse-autoencoder and expert-activation analyses (**Fig. 1g**) and scaled to human-proteome-scale library construction (**Fig. 1h**).

### OmniSyn achieves robust target-aware de novo generation with realistic chemical-space recovery without structural information

We first assessed whether OmniSyn had acquired a robust capacity for high-quality molecular generation in the de novo design task. As shown in **Fig. 2a**, OmniSyn achieved the highest normalized SA score (0.795), together with the highest atom-type (0.948), ring-type (0.912) and functional-group (0.752) distribution scores, while retaining competitive QED and diversity. Across a broader panel of physicochemical properties, including Fsp3, hydrogen-bond acceptor count, molecular weight, aromatic-ring count, rotatable-bond count and logP, OmniSyn most consistently reproduced the distributions of the reference active molecules **(Supplementary Figs. 4-5)**. In MolGenBench’s chemical-safety assessment, OmniSyn demonstrated strong chemical fidelity, yielding the highest scaffold-level chemical-filter score (0.445) and the second-highest SMILES-level score (0.604; **Fig. 2b**).

**Fig. 2.**
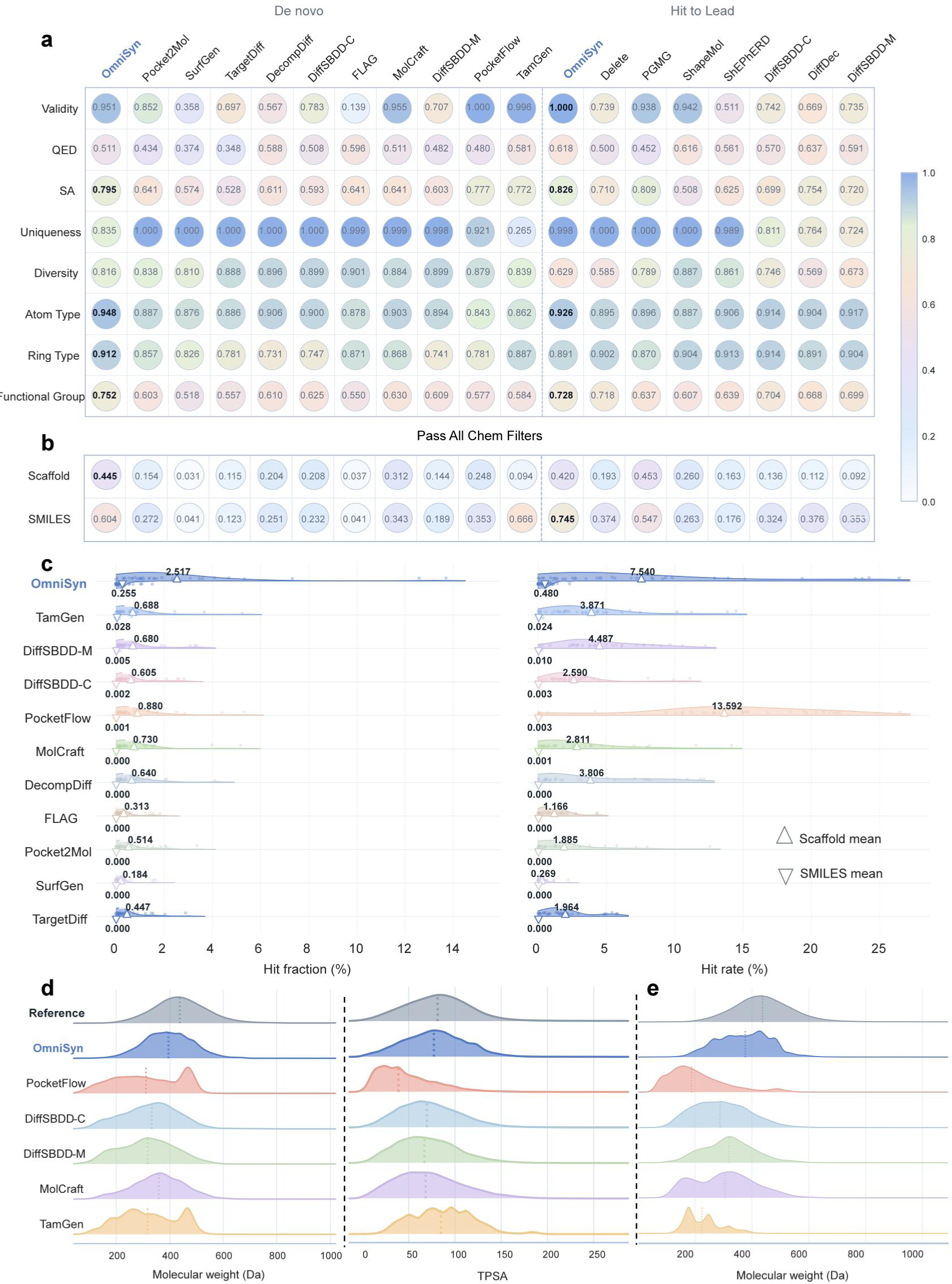
Chemical quality and active-molecule rediscovery in de novo generation. **a**, General molecular-property scores for de novo generation and H2L optimization. Metrics include validity, quantitative estimate of drug-likeness (QED), normalized synthetic accessibility (SA), uniqueness, diversity and agreement with reference atom-type, ring-type and functional-group distributions. Higher values indicate better performance; bold values denote the best result within each task. **b,** Fractions of generated scaffolds and complete molecular structures passing all MolGenBench chemical filters. **c,** Distributions of scaffold- and SMILES-level hit fractions (left) and hit rates (right) for the 35 unseen protein targets in MolGenBench. Each point represents the value obtained for one target. Upward- and downward-pointing triangles indicate the mean scaffold- and SMILES-level values across the 35 targets, respectively. **d,** Molecular-weight and topological polar surface area (TPSA) distributions of generated molecules compared with the reference active ligands. Vertical dotted lines indicate distribution means. **e,** Molecular-weight distributions of molecules contributing to scaffold-level rediscovery.

Generating chemically favorable molecules alone is not the ultimate goal of OmniSyn. We next tested whether, given only a protein sequence, the model could rediscover target-relevant active compounds. As shown in **Fig. 2c**, OmniSyn also recovered substantially more known active chemistry than baseline models. It achieved SMILES hit fraction and hit rate values of 0.255% and 0.480%, representing a **9.1-fold** and **20-fold** improvement, respectively, compared with 0.028% and 0.024% for the next-best SMILES-level baseline, TamGen. At the scaffold level, OmniSyn obtained the highest hit fraction (2.517%) and the second-highest hit rate (7.540%). We next compared the molecular-weight and TPSA distributions of molecules generated by the better-performing methods (**Fig. 2d**). OmniSyn closely matched the reference distributions, indicating that its target-aware generation preserves realistic physicochemical property profiles. We then focused on the recovered active molecules and found that several competing methods were strongly biased toward lower-molecular-weight compounds.Although PocketFlow^16^ produced a higher scaffold hit rate, its recovered scaffolds were predominantly associated with molecules having a mean molecular weight below 200 Da **(Fig. 2e)**. By contrast, OmniSyn scaffold hits followed a molecular-weight distribution closely resembled that of the reference actives, suggesting that its scaffold recovery reflects broader and more realistic active chemical space rather than enrichment for small, frequently occurring fragments. Together, these results show that OmniSyn combines strong target-aware generation with realistic physicochemical profiles and broad recovery of active chemical space, without relying on molecular-weight bias to improve scaffold recovery.

To further quantify OmniSyn’s target awareness that produce molecules with target-specific binding, we evaluated OmniSyn using the Target-Aware Score (TAScore) introduced in MolGenBench. At the SMILES level, more than 92% of targets fell within the lowest TAscore interval of 0-1 for most baselines, compared with only 26% for OmniSyn **(Fig. 3a)**. The remaining targets were shifted towards higher enrichment, with 60% and 14% in the 1-10 and 10-100 intervals, respectively. These results indicate that OmniSyn’s improved rediscovery performance is not simply driven by the generation of broadly active-like molecules, but instead reflects genuine target-specific conditioning from individual protein sequences.

**Fig. 3.**
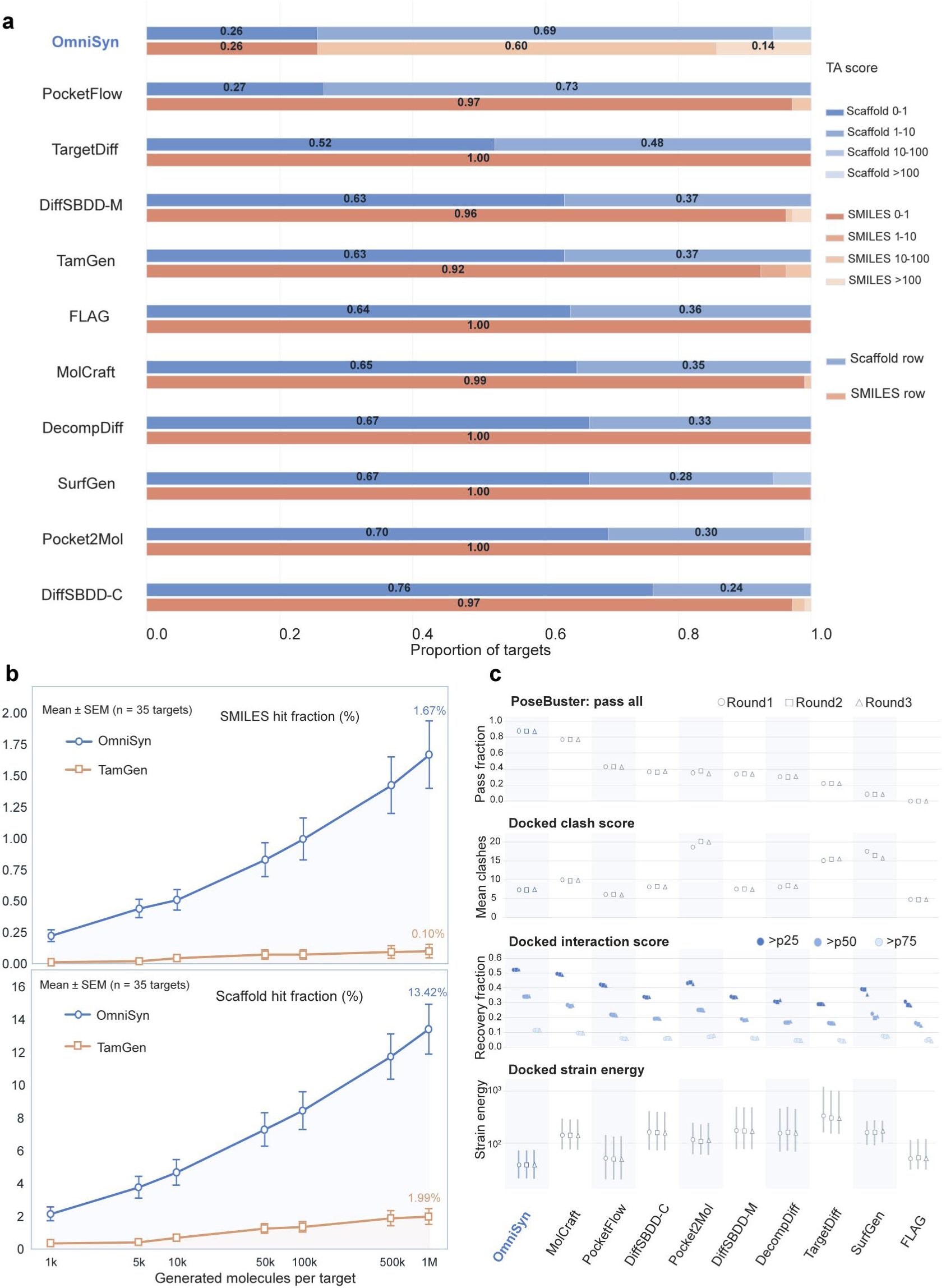
OmniSyn combines target-aware de novo generation with inference-time scaling and 3D plausibility. **a**, Distribution of scaffold- and SMILES-level TAScores across MolGenBench targets. Blue and orange bars denote scaffold- and SMILES-level evaluations, respectively, and colours indicate the proportions of targets within TAScore intervals of 0-1, 1-10, 10-100 and greater than 100. Higher intervals indicate stronger target-specific enrichment of recovered active molecules. **b,** Inference-time scaling of scaffold- and SMILES-level hit fractions as a function of the number of molecules generated per target. OmniSyn and TamGen were evaluated over matched sampling budgets ranging from 1,000 to 1 million molecules per target. Points show the mean across 35 targets, and error bars indicate the standard error of the mean (SEM). **c,** 3D evaluation following conformer generation and redocking. From top to bottom, panels show the PoseBusters pass-all fraction, mean docked clash score, recovery fractions at the benchmark p25, p50 and p75 docked-interaction thresholds, and docked strain energy. For strain energy, symbols indicate the median and vertical lines span the 25th-75th percentiles on a logarithmic scale. Circles, squares and triangles denote three independent evaluation rounds. Higher values are favorable for the PoseBusters and interaction-recovery metrics, whereas lower values are favorable for clash score and strain energy.

A fundamental challenge in de novo drug design is ensuring that models generalize to novel targets rather than relying on sequence homology from the training set. To assess OmniSyn’s robustness in this regard, we next tested whether its strong performance persisted under a stricter separation between training and evaluation proteins. OmniSyn was re-evaluated on a sequence-deduplicated MolGenBench subset defined using a 70% protein-sequence similarity threshold against the training data. This filtering criterion was stricter than that applied to the other evaluated models, creating a more challenging generalization setting. Rediscovery decreased modestly under this setting. Scaffold and SMILES hit fractions decreased from 2.517% to 2.027% and from 0.255% to 0.197%, respectively. The corresponding hit rates decreased from 7.540% to 5.516% and from 0.480% to 0.421%. Despite these reductions, OmniSyn retained the highest scaffold and SMILES hit fractions and the highest SMILES hit rate among the evaluated models (**Supplementary Fig. 6**). It also retained the highest SA (0.795), atom-type (0.947), ring-type (0.922) and functional-group (0.740) scores. OmniSyn further achieved the highest scaffold-level chemical-filter score (0.443) and the second-highest SMILES-level score (0.588). At the SMILES level, only 28% of sequence-deduplicated targets had TAScore below 1, compared with at least 92% for most baselines (**Supplementary Fig. 7**). OmniSyn also achieved the highest mean scaffold-level TAScore, with 64% of targets falling within the 1-10 interval and only 35% remaining in the lowest interval. Thus, stricter control of protein-sequence similarity caused only modest performance reductions and preserved OmniSyn’s comparative advantages across chemical quality, active-molecule recovery and target awareness.

### OmniSyn scales target-aware molecular recovery while preserving structural and synthetic feasibility

To fairly compare the performance against baseline models, our initial de novo benchmark evaluations used a sampling budget of 1,000 molecules per target. However, inspired by recent evidence of inference-time scaling in deep learning and large language models^43^, we asked whether molecular generation exhibits a similar dependence on inference budget. We therefore scaled sampling for OmniSyn and the strongest sequence-based baseline, TamGen^44^, from 1,000 to 1 million molecules per target and evaluated how target-relevant molecular recovery changed with increasing inference-time computation. As shown in **Fig. 3b**, OmniSyn showed sustained increases in both scaffold- and SMILES-level hit fractions, reaching 13.42% and 1.67%, respectively, at the largest budget. By comparison, TamGen reached only 1.99% at the scaffold level and 0.10% at the SMILES level with the same sampling budget. These results demonstrate favorable inference-time scaling in OmniSyn, whereby increased sampling computation consistently translates into broader recovery of target-relevant chemical space, with no clear saturation observed at 1 million molecules per target.

The favorable inference-time scaling of OmniSyn raises a further question: whether the target-relevant molecules also retain realistic 3D geometry and practical synthetic feasibility. We therefore subjected the generated molecules to independent structural and retrosynthetic evaluation. Following RDKit conformer generation and AutoDock Vina redocking^45^, OmniSyn achieved the highest PoseBusters^46^ pass-all fraction, approaching 0.9, and the strongest docked-interaction recovery across all evaluated percentile thresholds (**Fig. 3c**). Pass fractions for the individual PoseBusters validity, geometry and protein-ligand compatibility checks are reported in **Supplementary Fig. 8**. Its docked poses also exhibited low intermolecular clash scores and among the lowest strain energies. The absence of explicit pocket structures during generation therefore did not prevent OmniSyn molecules from adopting geometrically plausible, interaction-compatible poses under the MolGenBench evaluation protocol^35^.

Synthetic feasibility showed a similarly strong pattern. As shown in **Table 1**, OmniSyn achieved the highest AiZynthFinder^36^ success rate in the de novo setting (68.47%), outperforming the second-best method, TamGen, by 26.01%. This independent retrosynthesis assessment provides an external validation of the model-derived reaction traces: whereas OmniSyn jointly generates molecular structures and synthesis trajectories, AiZynthFinder evaluates their solvability using an independent retrosynthetic planner. Together, these results indicate that OmniSyn’s gains in target-aware recovery are accompanied by structurally plausible binding poses and independently supported synthetic feasibility, rather than arising from unconstrained expansion of chemical space.

**Table 1.**
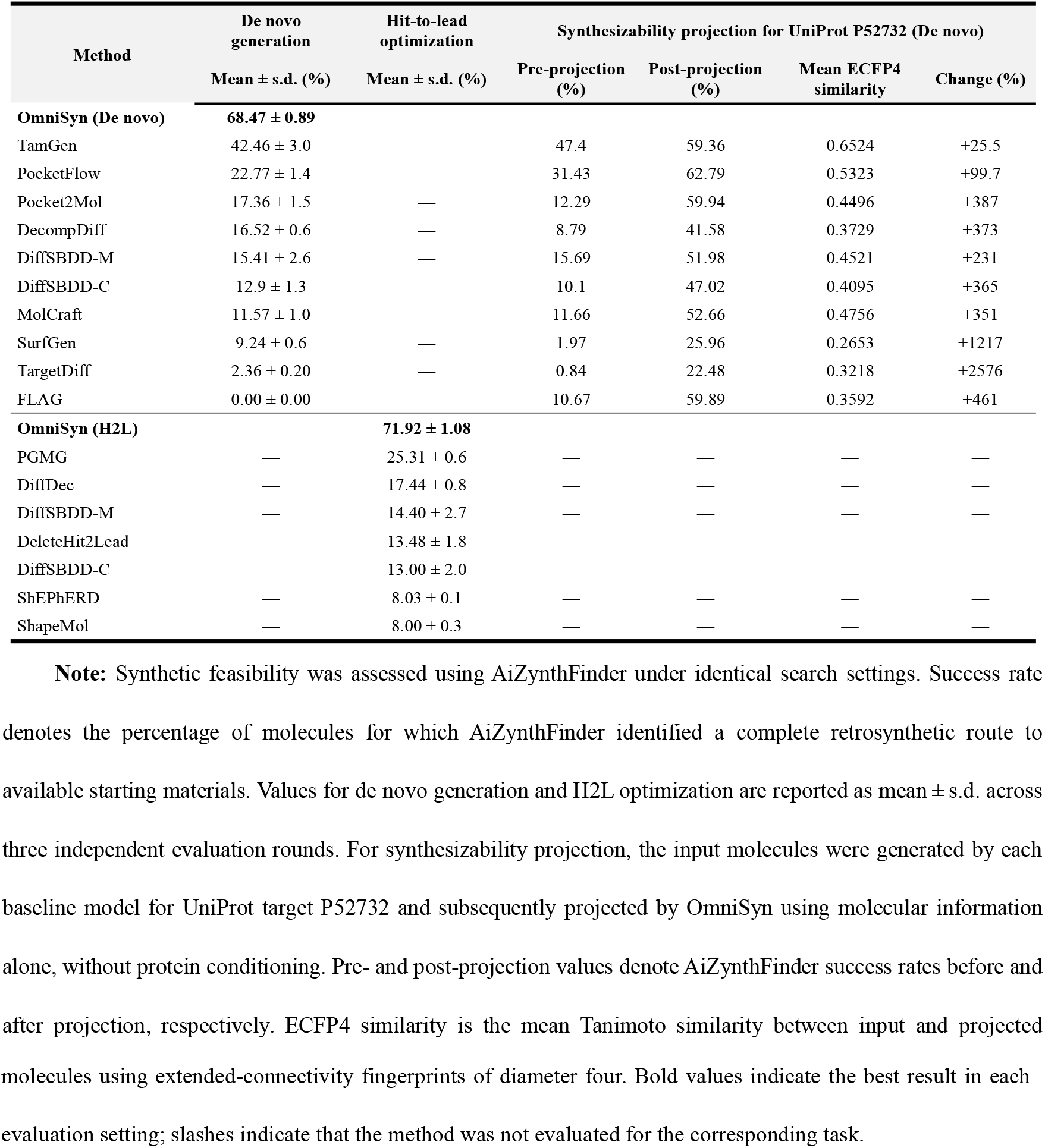
Independent AiZynthFinder evaluation of synthetic feasibility across OmniSyn design tasks.

### OmniSyn supports scaffold-guided H2L optimization with explicit synthesis routes

Alongside de novo design, H2L optimization constitutes another critical task of drug design that focuses on improving compound properties (e.g., binding affinity and selectivity) through partial atomic modifications, typically by exploring R-group variations while maintaining core scaffolds. We evaluated OmniSyn in the MolGenBench H2L task, in which generation is constrained by both a target and an existing molecular context. For each protein-scaffold pair, 10,000 candidate molecules were generated and ranked by their ECFP4 similarity to the input scaffold. The 200 most scaffold-similar candidates were retained for evaluation. In the property analysis presented alongside the de novo results, OmniSyn achieved the highest normalized SA score (0.826) and SMILES-level chemical-filter score (0.745), while maintaining complete validity and competitive QED (**Fig. 2a,b**). Its independently assessed AiZynthFinder success rate reached 71.92%, the highest among the evaluated models, outperforming the second-best method, PGMG, by 46.61 percentage points, supporting the synthetic tractability of the optimized molecules (**Table 1**). OmniSyn recovered active chemical series consistently across benchmark targets (**Fig. 4a**). It achieved the highest scaffold and SMILES hit rates, at 10.011% and 0.163%, respectively, while attaining scaffold and SMILES hit fractions of 9.025% and 0.986%. The simultaneous recovery of active scaffolds and exact molecular structures indicates that the model can preserve relevant chemical context while exploring productive modifications within established series. Consistent with this result, OmniSyn produced the largest number of SMILES-level hits while maintaining competitive mean normalized affinity (MNA; **Fig. 4b**), providing a favourable balance between recovery breadth and potency.

**Fig. 4.**
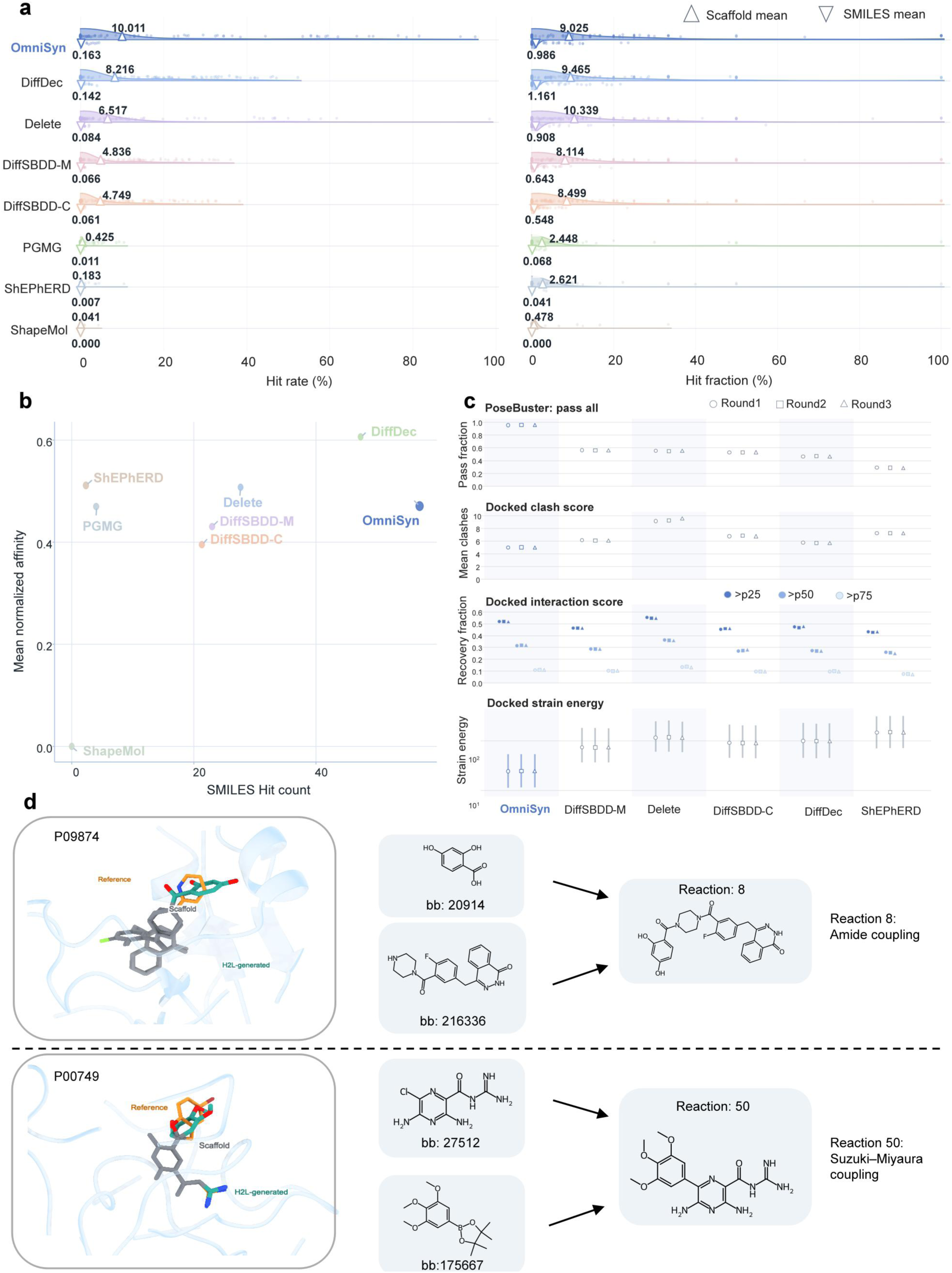
Target-aware and synthesis-traceable H2L optimization by OmniSyn. **a**, Distributions of scaffold- and SMILES-level hit rates (left) and hit fractions (right) across MolGenBench H2L targets. Upward and downward triangles denote the mean scaffold- and SMILES-level values, respectively. **b,** Relationship between the number of recovered SMILES-level hits and their mean normalized affinity (MNA). Models towards the upper right combine broader active-molecule recovery with stronger affinity. **c,** 3D evaluation of optimized molecules after conformer generation and redocking, comprising the PoseBusters pass-all fraction, mean docked clash score, interaction-score recovery above the 25th, 50th and 75th percentile thresholds, and docked strain energy. Circles, squares and triangles denote three independent rounds. **d,** Representative optimizations for targets P09874 and P00749. Binding-pose overlays show the reference scaffold and generated analogue; the accompanying schemes show the model-selected building blocks and reaction templates for amide coupling and Suzuki-Miyaura coupling, respectively.

Beyond recovering active chemical series, we further evaluated whether the optimized molecules retained plausible 3D compatibility with their targets after redocking (**Fig. 4c and Supplementary Fig. 8**). OmniSyn achieved the highest PoseBusters pass-all fraction and among the lowest docked clash and strain-energy values, together with consistently strong interaction-score recovery. Representative examples illustrate how this optimization is coupled to reaction-level traceability (**Fig. 4d**). For poly (ADP-ribose) polymerase 1 (PARP1; UniProt P09874), OmniSyn retained the reference scaffold while modifying its peripheral substituents through an amide coupling between a polyhydroxy benzoic acid building block and an amine-containing heterocyclic intermediate. For urokinase-type plasminogen activator (uPA; UniProt P00749), the retained heteroaryl scaffold was elaborated with a dimethoxyaryl group through a Suzuki-Miyaura coupling. In both cases, the generated analogue preserved the scaffold orientation observed in the reference binding pose and was returned with the corresponding building blocks and reaction template. These results show that OmniSyn connects target-aware lead optimization with explicit, model-derived synthesis routes. Together, these results establish OmniSyn as a sequence-conditioned H2L framework that jointly preserves scaffold context, target compatibility and synthetic tractability without requiring explicit pocket structures.

### OmniSyn converts synthetically inaccessible outputs from other generative models into close, synthesis-feasible analogues

While many external generative models produce molecules with desirable theoretical properties, these candidates frequently suffer from poor synthetic accessibility. To bridge this gap, OmniSyn further introduces a synthesizability projection capability, transforming molecules generated by external models into structurally related analogues with improved synthetic tractability. As shown in **Fig. 5a**, most projected molecules were predominantly shifted above the diagonal in normalized SA score while retaining high ECFP4 similarity to their inputs, indicating that improved accessibility was achieved through local structural modification rather than wholesale molecular replacement. This trend was independently reproduced by SCScore^47^, which decreased from a mean of 3.774 to 3.557, corresponding to a mean paired change of -0.216 (Wilcoxon test, = 8.9 × 10^−260^ ; **Fig. 5b**). Overall, 63.2% of projected molecules showed lower SCScore values, whereas only 32.0% increased, further demonstrating that OmniSyn can systematically redirect externally generated molecules toward more synthesis-accessible regions of chemical space while largely preserving their original molecular identity.

**Fig. 5.**
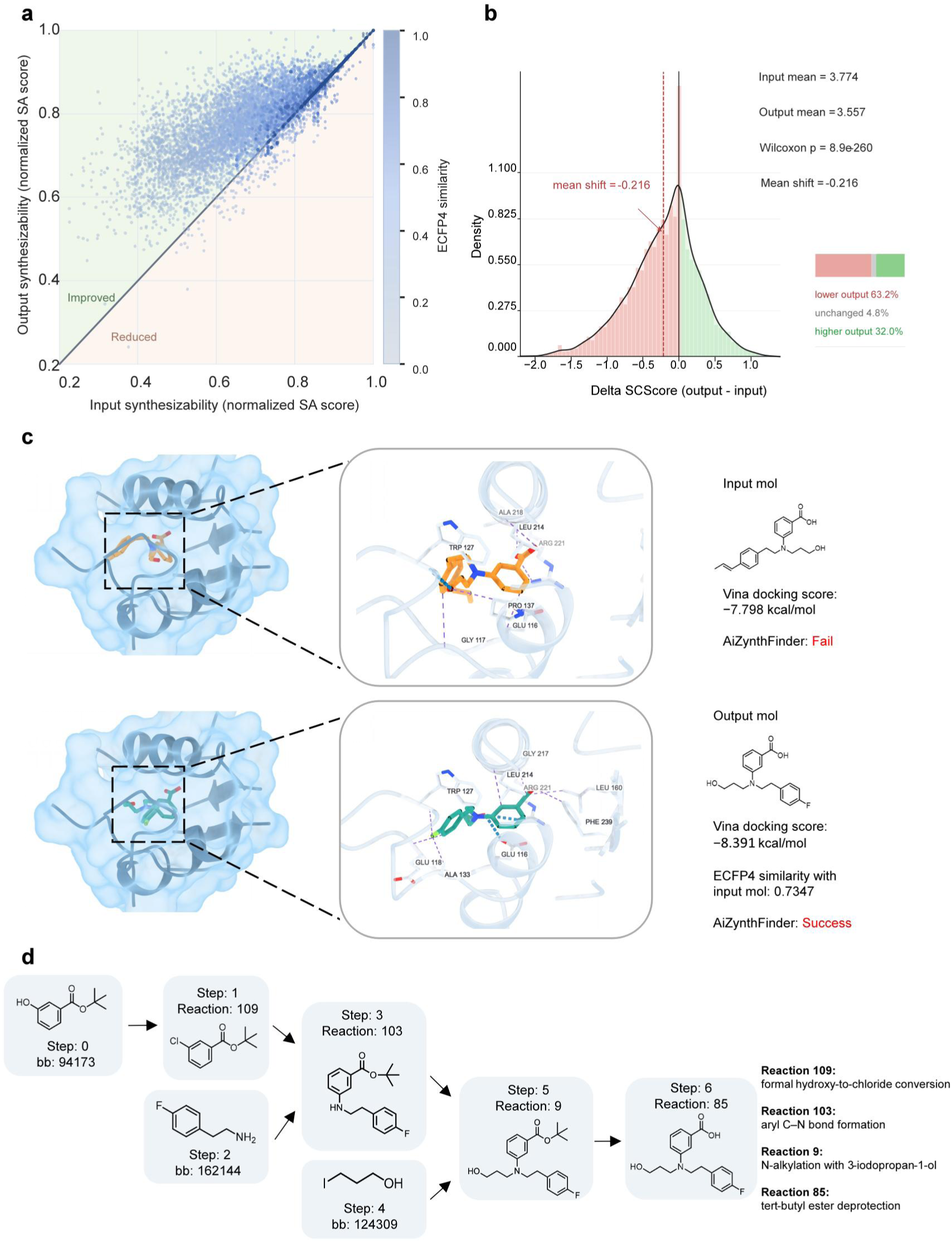
Synthesizability projection preserves molecular similarity while improving synthetic feasibility. **a**, Normalized synthetic-accessibility (SA) scores before and after OmniSyn projection. Points above the diagonal indicate improved synthetic accessibility, and colour denotes ECFP4 similarity between the input and projected molecules. **b,** Distribution of paired SCScore differences between projected and input molecules. Negative values indicate improved synthetic feasibility. Mean SCScore decreased from 3.774 to 3.557, with a mean change of −0.216 (Wilcoxon signed-rank test, = 8.9 × 10^−260^); 63.2% of molecules decreased, 4.8% remained unchanged and 32.0% increased. **c,** Representative input and projected molecules with their docked binding poses, Vina scores and AiZynthFinder outcomes. The projected analogue retained an ECFP4 similarity of 0.7347, improved the Vina score from −7.798 to −8.391 kcal/mol and changed the AiZynthFinder outcome from failure to success. **d,** Model-derived, template-level reaction route for the projected analogue.

A representative example illustrates how projection reconciled structural preservation with improved retrosynthetic feasibility (**Fig. 5c**). The input molecule failed AiZynthFinder analysis despite a Vina score of -7.798 kcal/mol. OmniSyn retained its tertiary-amine and benzoic-acid core while replacing the terminal substituent with a 4-fluorophenethyl group, yielding an ECFP4 similarity of 0.7347. The projected analogue preserved the principal polar-interaction region and extended the fluorophenyl substituent into an adjacent hydrophobic environment. It subsequently passed AiZynthFinder analysis and achieved a Vina score of -8.391 kcal/mol.

The projected molecule was generated together with a chemically interpretable, template-level route from purchasable building blocks (**Fig. 5d**). Starting from tert-butyl 3-hydroxybenzoate, Reaction 109 encoded a formal hydroxy-to-chloro conversion to give the corresponding aryl chloride. Reaction 103 then formed an aryl C–N bond with 4-fluorophenethylamine, yielding the protected secondary aniline while retaining the tert-butyl ester. Subsequent N-alkylation with 3-iodopropan-1-ol (Reaction 9) installed the 3-hydroxypropyl substituent. Finally, tert-butyl ester deprotection (Reaction 85) afforded the corresponding benzoic acid. This template-level route connects the structural changes introduced during projection to an explicit sequence of chemically recognizable transformations. A further representative projection example is provided in **Supplementary Fig. 9**. Together, these results establish synthesizability projection as a general synthesis-aware refinement strategy that redirects molecules from external generative models toward more accessible chemical space through minimal structural editing, while preserving their underlying molecular identity and target-relevant context.

### OmniSyn exhibits task and chemistry dependent specialization

The preceding benchmarks established OmniSyn’s performance across all three design settings, but did not reveal how its shared backbone adapts to their distinct conditioning requirements. We therefore interrogated OmniSyn across successive levels of model organization, from MoE expert routing to PLIM representations, pairwise attention and decoder states. Task-level routing profiles differed across PLIM blocks (**Supplementary Fig. 10**), showing that the three design settings recruited distinct combinations of experts, while retaining experts across related design settings. We next asked whether this computational allocation also varied with the chemical regime being processed. Across eight experts, routing frequencies showed distinct correlations with molecular weight, heavy-atom count, log*p*, ring count and the fraction of sp^3^-hybridized carbons (**Fig. 6a**). For example, expert *E*_0_ was preferentially used for smaller and less ring-rich molecules, with negative correlations with molecular weight (*r* =− 0.48), heavy-atom count (*r* =− 0.48) and ring count (*r* =− 0.39). By contrast, _6_ usage increased with heavy-atom count (*r* = 0.34) and ring count (*r* = 0.44), whereas _1_ was negatively associated with *F*_sp3_ (*r* =− 0.43). Together, these results show that PLIM routes computation according to both task identity and molecular regime, providing a network-level basis for multi-task adaptation.

**Fig. 6.**
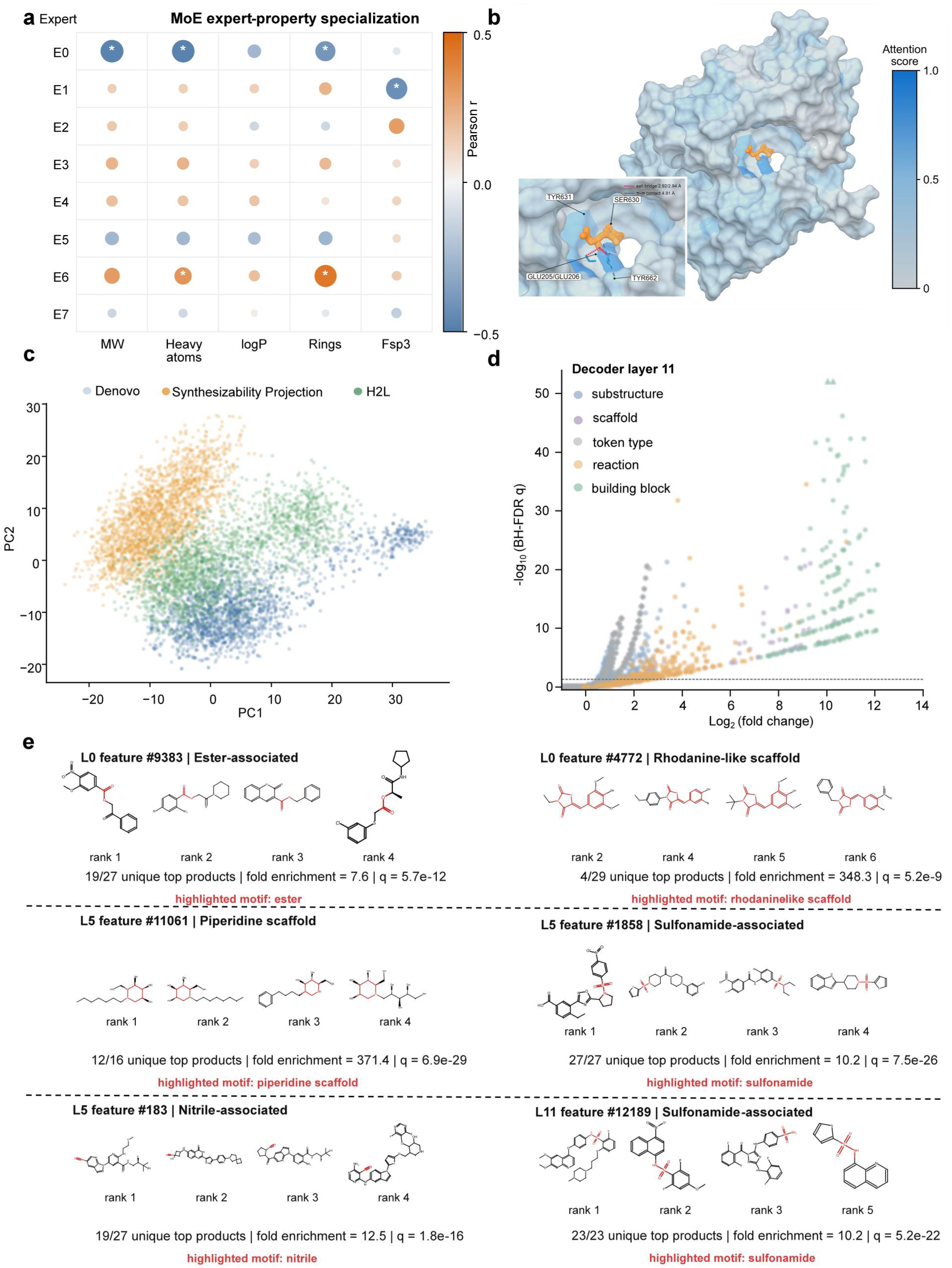
Task, interaction and chemical representations within OmniSyn. **a**, Pearson correlations between molecular descriptors and mean top-1 expert-routing frequencies across the four PLIM MoE layers, evaluated using 613 valid molecule-conditioned records representing 566 unique SMILES. These comprised 120 synthesizability-projection reference-molecule records and 493 H2L protein-scaffold records; protein-only de novo records were excluded because they contained no molecular input. Marker colour indicates Pearson’s *r*, marker size indicates −log(*q*), where *q* is the Benjamini-Hochberg-adjusted *P* value, and asterisks denote the six associations with the largest absolute correlation coefficients. **b,** Pair-attention analysis for a representative H2L example involving P27487. Sequence-derived attention scores were normalized and mapped post hoc onto the protein surface; the retained ligand scaffold is shown in orange. The inset highlights the scaffold-contacting region and neighboring residues Tyr631, Ser630, Trp659 and Tyr662. **c,** Principal-component projection of a fixed random sample of 6,000 PLIM conditioning representations drawn without replacement from a balanced pool of 300,000 stored activation records (100,000 per task). Points are colored by design task: de novo generation, synthesizability projection or H2L optimization. The projected representations were used as inputs to the conditioning SAE. **d,** Enrichment landscape for SAE features from decoder layer 11. Points show log_2_ fold enrichment against task-matched backgrounds and globally corrected significance for molecular substructures, scaffolds, token types, reaction-template identities and building-block identities. The horizontal line denotes a Benjamini-Hochberg false-discovery-rate threshold of *q* = 0.05 . **e,** Representative top-activating molecules for chemically associated SAE features from decoder layers 0, 5 and 11. Enriched motifs are highlighted in red. Statistics report the number of motif-containing molecules among unique valid top activations, log_2_ fold enrichment against the task-matched background and globally corrected values. SAE, sparse autoencoder; MoE, mixture of experts; MW, molecular weight; *F*_sp3_, fraction of sp3-hybridized carbon atoms.

Beyond the global adaptation achieved via expert routing, a robust model must also accurately perceive local protein-ligand interaction environments. Therefore, we next examined the finer-grained protein-molecule relationships captured by the PLIM. For a representative H2L example involving P27487, pair-attention scores were mapped onto the corresponding protein structure and retained ligand scaffold (**Fig. 6b**). The strongest attention was concentrated in the scaffold-contacting region of the binding site rather than distributed uniformly across the protein surface. Elevated scores occurred near Tyr631, Ser630, Trp659 and Tyr662, which surround the retained scaffold. Structural inspection further identified salt-bridge interactions involving Glu205 and Glu206, together with a π-π interaction involving Tyr662. Although attention scores are not physical interaction energies, their spatial correspondence with the binding environment indicates that PLIM localizes ligand-conditioned information to target regions relevant for scaffold-conditioned molecular optimization. This analysis extends the interpretability of OmniSyn from network-level expert allocation to residue-level protein-scaffold context.

To determine whether this interpretability extended from individual residue-scaffold relationships to global representation structure, we fitted TopK sparse autoencoders (SAEs; K=32)^47^ to the PLIM conditioning embeddings and to activations from decoder layers 0, 5 and 11. We first examined the sparse latent representations produced by the conditioning SAE. Principal-component analysis revealed a structured, task-dependent organization in which de novo generation, synthesizability projection and H2L optimization occupied distinct but partially overlapping regions (**Fig. 6c**). H2L representations were positioned between those of de novo generation and synthesizability projection. This organization mirrored the corresponding conditioning requirements: H2L optimization integrates both protein and molecular information, whereas de novo generation and synthesizability projection are driven primarily by protein and molecular inputs, respectively. The conditioning SAE therefore recovered a task-resolved representation space that preserved the three design settings while organizing them according to their input composition.

The conditioning SAE revealed global task organization, but we next asked whether individual sparse features were associated with final-product chemistry or synthesis-path decisions. Using task-matched feature and background populations, we tested the conditioning SAE and each decoder SAE for associations with molecular substructures, Bemis-Murcko scaffolds, token types, reaction-template identities and building-block identities. From the 4,096-feature conditioning dictionary and each 16,384-feature decoder dictionary, we selected 500 features before enrichment testing. Features were required to have an activation frequency of at least 10⁻⁴ and a task-specificity score of at least 1.5. Eligible features were ranked using a combined score defined as task specificity multiplied by the square root of activation frequency, and the top 500 were retained. Chemical enrichment tests used unique isomeric canonical final products as statistical units, whereas route tests used decoder states. One-sided hypergeometric *P* values were adjusted across all feature-attribute pairs within each representation using the Benjamini-Hochberg procedure. None of the 500 conditioning SAE features showed a significant chemical or route association after correction. By contrast, decoder SAE features exhibited numerous layer-resolved associations. Of the 500 features tested per layer, 48, 176 and 253 features at layers 0, 5 and 11, respectively, showed at least one significant chemical association. The corresponding numbers of features with route associations were 272, 457 and 450.

Chemical associations became increasingly prevalent with decoder depth, whereas route associations were abundant across all three layers and peaked at layer 5. The enrichment landscapes for layers 0 and 5 are shown in **Supplementary Fig. 11**, and that for layer 11 is shown in **Fig. 6d**. At layer 11, the strongest enrichments were dominated by building-block identities, with additional associations involving reaction templates and chemical information. Representative high-activation sets were enriched for final products containing ester, rhodanine-like, piperidine, nitrile and sulfonamide motifs (**Fig. 6e**). Additional chemistry- and route-associated examples across layers 0, 5 and 11 are shown in **Supplementary Figs. 12-14**. Together, these results reveal a multiscale internal organization. MoE routing specialized computation across tasks and molecular regimes, PLIM attention localized protein-scaffold context, and the conditioning SAE separated task states. Within the decoder, sparse features were associated with both final-product chemistry and specific building-block or reaction-template decisions.

### OmniSyn enables proteome-wide exploration of synthesis-accessible chemical space

A large fraction of the human proteome remains pharmacologically uncharted, limiting both the functional interrogation of disease-associated proteins and the development of therapeutics beyond established target classes. Systematically designing target-specific, synthesizable molecules could therefore expand the accessible chemical space of human proteins and provide starting points for chemical probes and candidate inhibitors. Having established OmniSyn’s target-aware molecular design independent of protein crystal structures, we next investigated whether conditional generation could be scaled from benchmark targets to the human proteome. This transition requires high sampling throughput because a distinct candidate set must be generated for each protein sequence. Within 120 minutes, OmniSyn sampled 2.12 million and 1.92 million molecules in the de novo and H2L tasks, respectively (**Fig. 7a**). Its de novo sampling rate was 3.2-fold higher than that of TamGen, whereas its H2L sampling rate was approximately two orders of magnitude higher than those of the evaluated 3D baselines. This throughput provided the basis for scaling sequence-conditioned generation to the human proteome. To pursue this goal at proteome scale, we applied OmniSyn to 21,306 human protein sequences, generating 2.7 billion target-molecule entries with explicit synthesis traces. The candidates were scored using our previously developed PSICHIC-plus model and Vina docking and then filtered to construct a searchable library for computational prioritization and automated experimental follow-up (**Fig. 7b**). To our knowledge, this represents the largest human-proteome-scale generative molecular library reported to date. For comparison, DrugCLIP^37^ screened a fixed collection of 500 million compounds against approximately 10,000 proteins, whereas the OmniSyn library covered more than twice as many targets and contained 5.4-fold more molecular entries (**Fig. 7c**). DrugCLIP ranks molecules from a shared predefined collection, whereas OmniSyn conditionally generates a distinct, synthesis-traceable candidate set for each protein sequence.

**Fig. 7.**
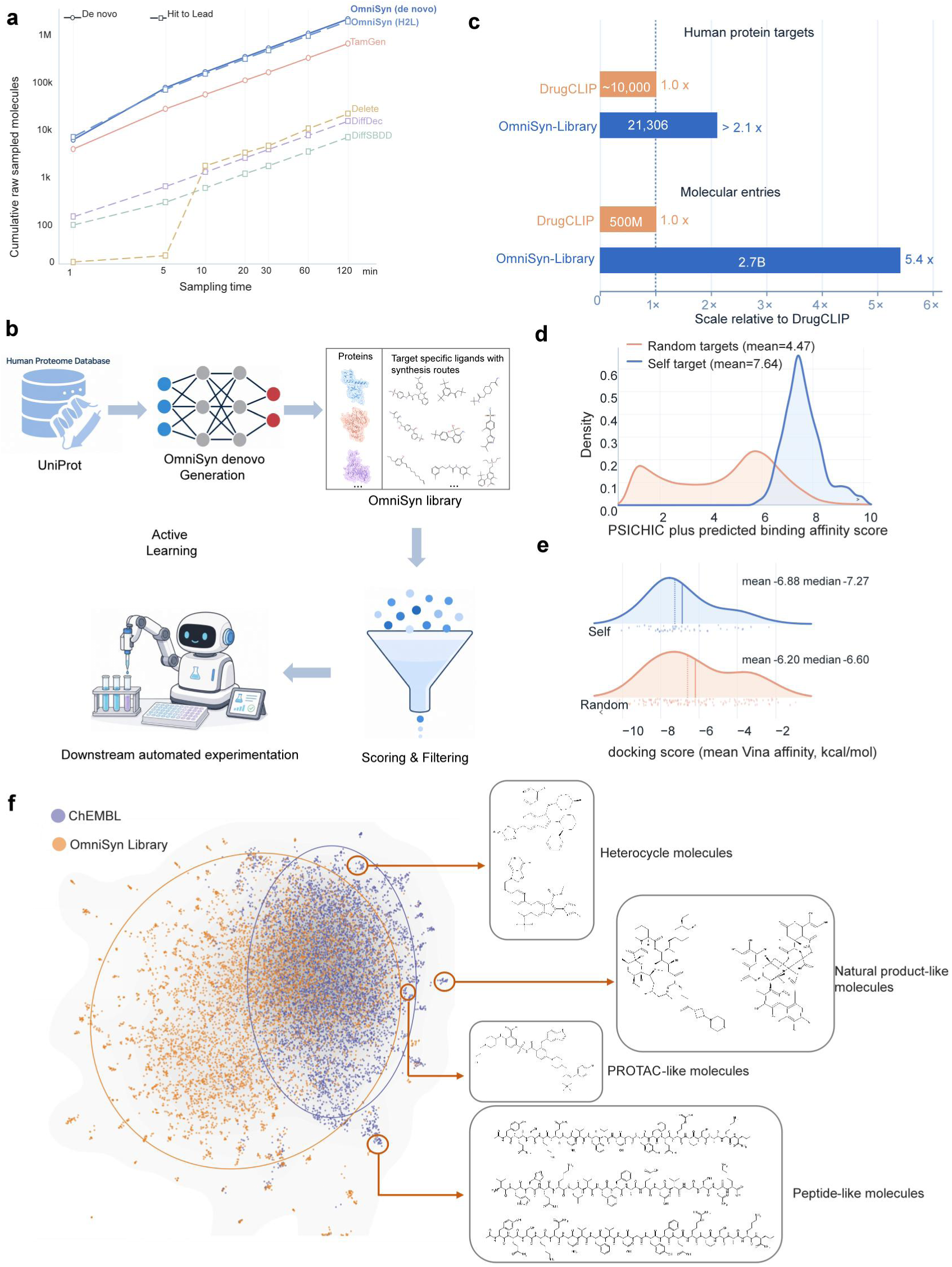
Proteome-scale construction and computational characterization of the OmniSyn molecular library. **a**, Sampling throughput for de novo and H2L generation. OmniSyn produced raw molecules faster than the baseline methods across the measured time range. Both axes are logarithmically scaled; 1-min values were interpolated or estimated, whereas subsequent values were recorded directly. **b,** Library-construction workflow. Human protein sequences from UniProt conditioned OmniSyn generation, followed by scoring, filtering, automated experimental follow-up and active learning. **c,** Scale relative to DrugCLIP. OmniSyn covers 21,306 human proteins and 2.7 billion target-specific molecular entries, corresponding to more than 2.1 times as many targets and 5.4 times more entries. **d,** PSICHIC-plus affinity-score distributions for molecules paired with their conditioning proteins (self-targets; 10,000 pairs from 200 targets) or non-cognate proteins (random targets; 99,300 pairs). Higher scores indicate stronger predicted interactions. **e,** Mean AutoDock Vina score distributions for self (n = 48) and random (n = 136) protein–molecule pairs. More negative scores indicate more favourable predicted binding; vertical lines denote the mean and median. **f,** Joint two-dimensional chemical-space embedding of OmniSyn-library and ChEMBL-derived representatives. Highlighted regions more extensively represented by ChEMBL contain structurally complex heterocyclic, natural-product-like, PROTAC-like and peptide-like molecules.

Scale alone, however, does not establish whether generated molecules retain a preference for their conditioning targets. For each of 200 proteins randomly selected from the human-proteome library, generated molecules were ranked using PSICHIC-plus (an enhanced version of PSICHIC^48^) according to their predicted scores for the conditioning protein. The top 50 candidates for each protein were retained and cross-scored against randomly selected non-cognate proteins using the same model. The candidates received higher scores for their conditioning proteins than for the non-cognate proteins (mean, 7.64 vs 4.47; **Fig. 7d**). Because PSICHIC-plus was used for candidate selection, we next examined whether this preference persisted under an orthogonal structure-based assessment. Vina docking likewise assigned more favorable scores to the conditioning proteins than to the non-cognate proteins (mean, - 6.88 vs -6.20 kcal/mol; **Fig. 7e**). Together, the sequence- and structure-based assessments support target-conditioned enrichment among the prioritized library candidates.

Target-conditioned enrichment could nevertheless arise from reproducing target-associated training molecules rather than exploring related active chemical space. We therefore examined this possibility through a case study of programmed death-ligand 1 (PD-L1), which is an immune-checkpoint ligand that attenuates antitumor T-cell responses and contributes to tumor immune evasion^49^. Generated candidates retained after filtering with PSICHIC-plus score, Vina docking score^45^ and DrugCLIP score^34^ were compared with PD-L1-associated training molecules and a held-out reference set of known PD-L1 actives. The reference set excluded all molecules present in the OmniSyn training data. The similarity distribution lay predominantly above the diagonal, indicating that the generated candidates were generally more similar to held-out actives than to their nearest training molecules (**Supplementary Fig. 15**). Two representative candidates showed similarities of 0.66 and 0.62 to held-out actives, compared with 0.39 and 0.36 to their closest training molecules, respectively. These results argue against simple nearest-neighbour memorization and indicate that OmniSyn recovers active-like chemical space beyond that represented by the target-associated training molecules.

Beyond the PD-L1 case study, the value of a proteome-scale library also depends on its global coverage of bioactive chemical space. To compare this coverage at tractable scale, we constructed diversity-stratified samples of 50,000 OmniSyn-library molecules and 50,000 ChEMBL 36-derived molecules and jointly embedded their ECFP4 fingerprints using UMAP^50^ (**Methods**). The resulting two-dimensional embedding showed extensive overlap between the two sets, together with substantial regions preferentially occupied by generated molecules (**Fig. 7f**). OmniSyn generation therefore extended beyond the chemical regions most extensively shared with the ChEMBL-derived reference set. Regions more extensively represented by ChEMBL contained peptide-like, PROTAC-like, structurally complex heterocyclic and natural-product-like molecules, highlighting complementary chemical regimes beyond those most readily accessed by the current building-block and reaction-template vocabulary. Collectively, these results establish OmniSyn as a scalable platform for constructing proteome-wide, target-specific and synthesis-traceable molecular resources.

## Discussion

The dominant paradigm in target-aware molecular design treats protein structure as the conditional signal and synthesizability as a property to be filtered for after the fact^14–16^. OmniSyn demonstrates that these assumptions are not the only viable route for scalable molecular design. Conditioned only on protein sequence, it exceeded a broad panel of structure-based generators on active-molecule rediscovery and target awareness^35^ while associating each generated molecule with a synthesis trace. The deeper implication is a reframing of the problem itself. Target relevance and synthetic accessibility, traditionally treated as objectives optimized sequentially, are here recovered as a joint optimization problem: by making synthesis actions part of the generative space rather than a downstream filter^23–26^, OmniSyn constrains exploration toward molecules compatible with its learned synthesis space. What is usually a late-stage triage step becomes a constraint integrated into generation, and the explored chemical space is therefore aligned with experimentally actionable chemistry.

That a sequence-conditioned model can achieve competitive target conditioning without explicit structural inputs provides insight into where useful biological information may reside. Structure has often been treated as the necessary intermediary between a protein and its ligands, but predicted structures represent one approximation of sequence-encoded biochemical constraints, and errors introduced during structure prediction can propagate into downstream design. By conditioning directly on sequence, OmniSyn avoids dependence on an explicit structural prediction step and its associated uncertainties. The interpretability analyses reinforce this reading: attention concentrated on scaffold-contacting residues; expert routing that separated design tasks and tracked molecular regime, and decoder features aligned to recognizable medicinal-chemistry motifs. Together, these observations indicate that OmniSyn learns structured internal representations of protein context, molecular state and synthesis chemistry, rather than relying on task-specific shortcuts.

These properties also expand what a generative model can accomplish. By sampling synthesis-traceable molecules directly for each target sequence rather than ranking fixed libraries, OmniSyn enables the design space to scale with target diversity rather than library size, shifting the unit of discovery from the library to the target. The synthesis-native conditioning also enables OmniSyn to jointly optimize multiple molecular design regimes within a single framework. Rather than treating de novo generation, synthesizability projection and H2L optimization as independent tasks, OmniSyn learns shared representations across these regimes, allowing knowledge transfer between complementary objectives. This multi-task synergy improves the model’s ability to capture target context, molecular transformation patterns and synthesis constraints, thereby enhancing performance and generalization across individual design tasks.

Despite these advances, OmniSyn’s accessible chemical space remains constrained by its reaction vocabulary and building-block inventory, which limits exploration beyond currently reachable synthesis space, particularly for scaffold-preserving H2L optimization. Expanding reaction diversity and molecular editing capabilities will further broaden this space. Moreover, a chemically valid synthesis trace does not guarantee experimental success, as practical synthesis depends on factors such as reaction conditions, yields and purification. Incorporating experimentally informed synthesis feasibility models into future optimization frameworks will be an important next step.

Taken together, these results suggest a shift in AI-driven molecular design, in which diverse design objectives are optimized within a unified generative process rather than addressed through sequential filtering. The central advance of OmniSyn is therefore not simply improved molecular generation, but the integration of biological context, molecular state and synthesis actions within a single adaptive learning framework. This integration provides a foundation for an active-learning design-make-test loop that couples sequence-conditioned generation and retrosynthetic reasoning to automated synthesis and biological assays (**Fig. 7b**). In each cycle, OmniSyn could propose synthesis-traceable candidates, automated experiments could evaluate synthetic execution and target activity, and the resulting outcomes could be returned to the model to guide subsequent rounds of generation. Such feedback could progressively align the model’s learned chemical space with experimentally accessible and target-relevant chemistry, while reducing its reliance on static historical datasets. Realizing this vision will require prospective validation of molecular activity, synthetic execution and closed-loop model updating, but OmniSyn provides a step towards transforming AI-designed molecules from computationally plausible structures into experimentally actionable candidates.

## Acknowledgements

This work was supported by the National Natural Science Foundation of China (Grant Nos. 82373937 to J.Y. and U25A20568 to F.K.), the Fundamental Research Funds for the Central Universities (Grant No. 22120260486), and the Zhongguancun Academy, Beijing 100094, China (Grant No. XTS0047). X.H. was supported by the National Natural Science Foundation of China (Grant Nos. 22633005, 92477103 and 22273023), the Shanghai Municipal Science and Technology Commission (Grant No. 25511102400), the Shanghai Municipal Commission of Economy and Informatization (Grant No. 2026-GZL-RGZN-01015), the Advanced Materials–National Science and Technology Major Project (Grant No. 2026ZD0623702), the Shanghai Frontiers Science Center of Molecule Intelligent Syntheses, and the Fundamental Research Funds for the Central Universities. We acknowledge the Bioinformatics Supercomputing Center at the School of Life Sciences and Technology, Tongji University, and the Supercomputer Center of East China Normal University (ECNU Multifunctional Platform for Innovation 001) for providing computational resources and support.

## Author Contributions Statement

Z.Q., Y.L., Y.Z., Y.Z. and H.Y.K. contributed equally. D.C., X.H., J.Y., Y.M., and Z.W. conceived and supervised the research project. Z.Q. and Y.L developed the primary method and code, and assisted in the analysis of the primary baselines and data. Z.Q., Y.L. and D.C. wrote the paper. All authors read and approved the manuscript.

## Methods

### Design rationale and task definition

OmniSyn was designed to make a reaction-based generator responsive to protein, molecular and task context without sacrificing its learned synthesis prior. The model therefore had to connect three heterogeneous design settings to the same decoder. We separated this problem into conditional representation learning, decoder alignment and task-level optimization. PLIM integrates protein and molecular information with task identity and allocates conditional capacity through sparse experts. Route self-distillation teaches the decoder to interpret these representations using action-level supervision from high-quality pseudo-trajectories. GDPO then optimizes task-specific molecular objectives while keeping the adapted decoder fixed. This staged design supports protein-conditioned de novo generation, molecule-conditioned synthesizability projection and protein- and scaffold-conditioned H2L optimization within a single synthesis-traceable framework.

Let *P* denote a protein sequence, *m*_0_ an input molecule or scaffold when present, *m* the generated product and *k* ∈ {1,2,3} the task identity. Task 1 maps *P* to *m* for protein-conditioned de novo generation. Task 2 maps *m*_0_ to for molecule-only synthesizability projection and never receives protein information. Task 3 maps (*P*, *m*_0_) to *m* for protein- and scaffold-conditioned H2L optimization; *m*_0_ is an input condition and is not forced into the decoder prefix. In every task, the decoder emits a sequence of synthesis actions *τ* = (*a*_1_, …, a*_τ_*), including token-type, building-block and reaction-template decisions, and a completed valid trajectory is deterministically decoded to a final molecule and an explicit synthesis trace.

### Task-specific dataset construction

Training datasets were constructed separately for the three conditional tasks from Papyrus and ChEMBL 36, with the latter providing measurements more recent than those incorporated into the Papyrus release. Because the three tasks require different biological and chemical contexts, no single activity-type or potency criterion was applied uniformly across them. Molecular structures were standardized and deduplicated, and records requiring a protein condition were mapped to representative protein sequences. Records associated with the 35 MolGenBench evaluation unseen targets were excluded from the primary training sets to reduce direct benchmark overlap.

Task 1 comprised curated positive protein-ligand bioactivity records for protein-conditioned de novo generation. Task 2 pooled eligible molecules into a protein-free corpus for molecule-conditioned synthesizability projection. Task 3 organized target-associated compounds into experimentally annotated or chemically inferred series, extracted a representative maximum common substructure for each series, and paired the protein sequence and series context with individual member molecules for H2L optimization.

In addition to exact benchmark-target exclusion, sequence-deduplicated counterparts were constructed by removing training proteins sharing at least 70% sequence similarity with any MolGenBench target using MMseqs2. Detailed source-specific inclusion criteria, molecular standardization, deduplication, chemical-series construction, dataset sizes and physicochemical distributions are provided in the **Supplementary Figs. 1-3**.

### Inference

The task is selected entirely through the input condition and task token. Task 1 receives a protein sequence; Task 2 receives a seed molecule only; and Task 3 receives a protein sequence plus molecular scaffold or fragment. The frozen OmniSyn decoder samples trajectories autoregressively from the PLIM-derived condition. Valid completed trajectories are decoded to products and stored with their building-block and reaction-template traces.

### MolGenBench evaluation

For de novo evaluation, OmniSyn generated 1,000 molecules for each of the 35 unseen protein targets in MolGenBench. For H2L evaluation, 10,000 candidates were generated for each protein-scaffold pair. These candidates were ranked by their ECFP4 Tanimoto similarity to the input scaffold, and the top 200 were retained for subsequent evaluation.

General molecular properties and target-recovery performance were evaluated following the MolGenBench protocol, using the benchmark-released outputs for baseline comparison. The evaluated properties included validity, QED, normalized synthetic accessibility (SA), uniqueness, diversity, and agreement with reference atom-type, ring-type and functional-group distributions. Higher normalized SA values indicate greater synthetic accessibility, while ChemFilter measures the fractions of scaffolds and complete molecules passing medicinal-chemistry filters. Scaffold- and exact-SMILES-level hit rates quantify active chemistry within the generated set, whereas hit fractions measure the coverage of target-associated reference actives. The Target-Aware (TA) score evaluates the target specificity of the recovered chemistry, with higher values indicating stronger target-specific enrichment.

For both de novo and H2L evaluation, molecules included in the benchmark analysis were converted into three-dimensional conformers and docked to their corresponding protein targets using AutoDock Vina. The resulting poses were then evaluated following the MolGenBench protocol, including PoseBusters pass-all fraction, intermolecular clash score, docked strain energy and interaction-score recovery.

### AiZynthFinder evaluation

AiZynthFinder was used as an external retrosynthetic planner under identical search-policy and stock settings for all compared models. A molecule was counted as solved when the planner returned a complete route to available stock materials. For each de novo and H2L model, up to 10,000 RDKit-valid molecules were selected across the three MolGenBench rounds (3,333, 3,333 and 3,334 candidates). Canonical SMILES were unique across rounds within a model. No molecular-weight window was applied.

To balance coarse chemical-property regimes, candidates were binned jointly by molecular-weight interval floor (MW/100), rotatable-bond bin min (floor(*N*_rot_/3), 3) and aromatic-ring bin min (*N*_arom_, 3). Molecules within bins were shuffled and bins were traversed in deterministic round-robin order with seed ‘ 42:task:model ’ . If a round contained fewer eligible unique molecules than requested, all available molecules were retained.

For synthesizability projection, baseline-generated molecules for MolGenBench target P52732 were supplied to OmniSyn in molecule-only Task 2 mode. The original and projected molecules were compared by molecular similarity and AiZynthFinder solution status.

### Sparse-autoencoder analysis

SAE analysis used the final checkpoint. This checkpoint contains PLIM and the decoder whose Transformer layers 0, 10 and 11 were adapted during self-distillation; during GDPO, the decoder was frozen and only PLIM was updated. Activations were collected in evaluation mode with gradients disabled. SAE-Cond used the 1,024-dimensional PLIM conditioning vector. SAE-Dec used token-aligned hidden states from decoder layers 0, 5 and 11, obtained by teacher-forcing offline trajectories sampled on-policy by the final model; the state at position was aligned with the action predicted at *t*+ 1.

TopK sparse autoencoders were trained with *K* = 32 and mean-squared reconstruction error, without an *l*_1_ penalty. The conditioning SAE used a fourfold expansion (4,096 features), and each decoder SAE used a sixteenfold expansion (16,384 features). Decoder dictionary vectors were normalized during training and the decoder bias was initialized from the empirical activation mean.

For each feature, we recorded activation frequency, mean activation and per-task activation frequency, and retrieved the top-activating samples with their task, molecular and route metadata. One-sided hypergeometric enrichment tests with Benjamini-Hochberg correction were applied only to discrete annotations: molecular substructures, Murcko scaffolds, token type, building-block identity and reaction-template index. Molecular weight and QED were summarized descriptively as mean values among top-activating samples versus task-matched background samples; they were not subjected to the hypergeometric test.

### Human-proteome virtual-library generation

The archived generation set comprised 21,306 canonical human protein sequences from UniProt. Each sequence was supplied independently in Task 1 mode, and every stored candidate record linked the target identifier, generated product and model-derived synthesis trace. The reported library size of 2.7 billion denotes target-molecule entries: the same chemical structure conditioned on different targets contributes distinct entries and the value should not be interpreted as a count of globally unique molecular structures. Candidate scoring and filtering were applied downstream for prioritization, including the PSICHIC-plus ranking and orthogonal docking analyses described in the main text.

### Chemical-space comparison with ChEMBL

Chemical-space coverage was compared using hierarchical diversity sampling of the OmniSyn library and a ChEMBL 36-derived reference set. For OmniSyn, molecules generated across all sampling rounds were pooled separately for each protein and deduplicated by SMILES. RDKit-parsed molecules were represented by 2,048-bit radius-2 Morgan fingerprints (ECFP4). Up to 150 molecules per protein were retained by MaxMin diversity selection using Tanimoto distance (seed 42). These per-protein representatives were pooled, assigned to 2,000 MiniBatchKMeans^51^ clusters and sampled to 50,000 molecules using cluster quotas proportional to the square root of cluster size.

For the ChEMBL reference set, unique canonical SMILES were represented using the same ECFP4 definition, assigned to 1,000 MiniBatchKMeans clusters and sampled to 50,000 molecules using the same cluster-weighting procedure. MiniBatchKMeans used a batch size of 10,000, three initializations, a maximum of 100 iterations and a random seed of 42. The two representative sets were jointly embedded using UMAP with Jaccard distance (number of neighbors = 15, minimum distance = 0.6, spread = 1.5, PCA initialization and random seed 42). All 100,000 embedded molecules were plotted. A pooled kernel-density estimate and separate two-standard-deviation covariance ellipses were included for visualization. Plot limits were determined from the 1st and 99th percentiles of each UMAP coordinate with 10% padding.

### Data availability

The data presented in this study are derived exclusively from publicly available datasets. The data for MolGenBench evaluation is available at https://zenodo.org/records/18 <u>183463</u>.

### Code availability

The code used to generate the results reported in this study will be made publicly available upon acceptance at the following GitHub repository: https://github.com/Intelligent-Drug-Discovery-Lab/OmniSyn.

