## Supplementary Information for "OmniSyn unifies target-aware molecular generation and optimization within a synthesis-native LLM framework across the human proteome"

**Table of Contents**

**S1 Supplementary Information of OmniSyn training data**

**S2 Supplementary Information of physicochemical distributions generated by different models**

**S3 Supplementary Information for MolGenBench evaluation results after deduplicating by similarity**

**S4 Supplementary Information for detailed PoseBusters evaluation of generated molecules**

**S5 Supplementary Information for Additional example of synthesizability projection by OmniSyn**

**S6 Supplementary Information for Task-dependent expert routing with PLIM**

**S7 Supplementary Information for SAE analysis**

**S8 Supplementary Information for case study in PD-L1**

**S9 References**

### Supplementary Information of OmniSyn training data

#### Task 1: protein-conditioned de novo generation

The Task 1 dataset was constructed by integrating bioactivity records from Papyrus1 and ChEMBL 36[2,3]. Because the Papyrus release used here incorporates data from ChEMBL 30, ChEMBL 36 was included to capture more recent measurements. Papyrus records were mapped to canonical SMILES and representative protein sequences. We retained positive quantitative measurements reported as Ki, Kd, IC50 and EC50 or pIC50, together with categorical records annotated as binders. Inactive records and synthetic negative examples were excluded. For ChEMBL 36, compound-target records were retained when the reported activity was below 10 μM.

After merging the two sources, records were deduplicated at the compound-target level. To limit the dominance of extensively characterized proteins, molecules associated with targets containing more than 100 active compounds were clustered using ECFP4 fingerprints at a Tanimoto similarity threshold of 0.6. One representative was retained from each cluster, selected using the strongest available activity measurement. This procedure reduced redundancy within individual targets while preserving distinct active chemical series.

We constructed two versions of the Task 1 training corpus with different levels of separation from the 35 MolGenBench targets (**Supplementary Fig. 1**). In the primary setting, records associated with the exact benchmark target identities were removed. For the more stringent setting, MMseqs2 was used to compare every training protein with the benchmark target sequences, and training proteins sharing at least 70% sequence similarity with any benchmark target were excluded. The exact-target-removal dataset contained approximately 0.60 million activity records, 0.42 million compound entries and 0.17 million unique molecules. The 70% sequence-deduplicated corpus retained approximately 0.59 million records, 0.41 million compound entries and 0.17 million unique molecules.

Sequence-level deduplication had little effect on the composition of the resulting chemical corpus. Binder annotations remained the largest activity category, contributing approximately 0.23 million records, followed by pIC50 and IC50 measurements at approximately 0.15-0.16 million and 0.14 million records, respectively. The molecular-property distributions were likewise preserved: the mean molecular weight changed only from 431.34 to 431.31 Da, whereas the mean QED remained 0.51 in both settings. Thus, the stricter sequence-based exclusion reduced potential target-level leakage without substantially altering the scale, activity composition or drug-like chemical characteristics of the Task 1 dataset.

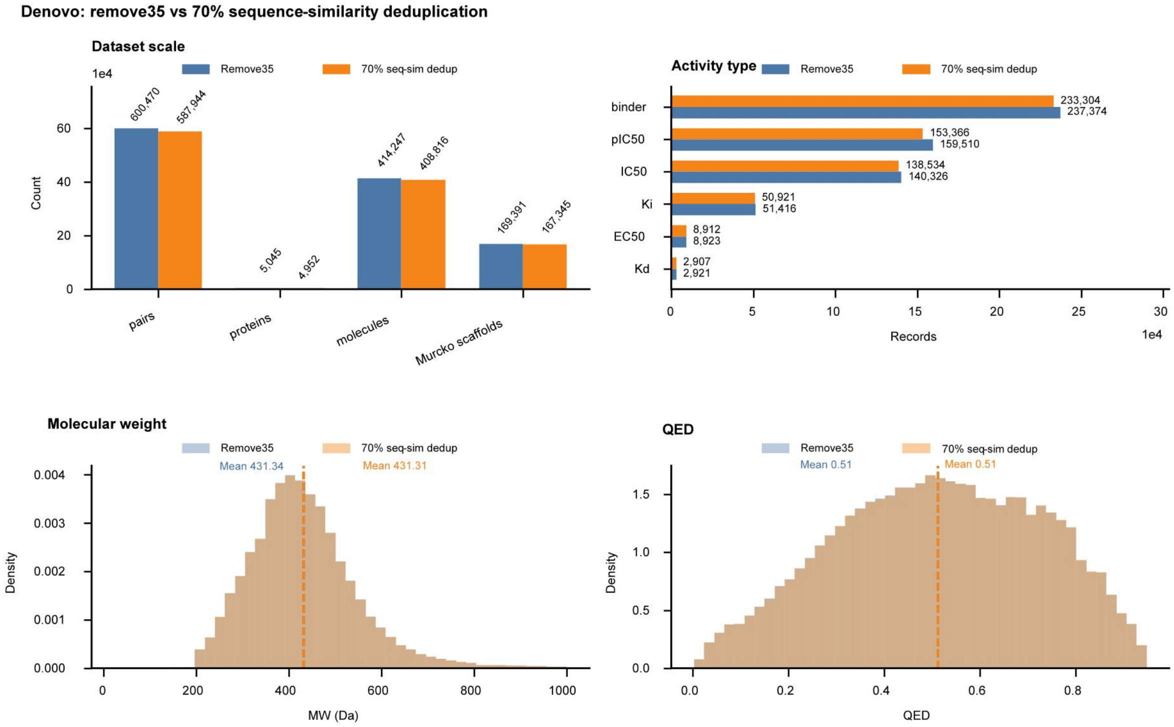


**Supplementary Fig. 1 | Construction and sequence-level deduplication of the Task 1 dataset. a,** Dataset scale after removing records associated with the exact 35 MolGenBench targets and after applying 70% protein-sequence-similarity deduplication with MMseqs2. Bars show the numbers of bioactivity records, protein targets, compound entries and unique molecules. **b,** Numbers of retained records for each activity type, including categorical binder annotations and quantitative $K_i$, $K_d$, $IC_{50}$ and $EC_{50}$ or $pIC_{50}$ measurements. **c,d,** Molecular-weight (**c**) and QED (**d**) distributions of molecules retained under the two deduplication settings. Dashed vertical lines and accompanying values indicate distribution means. Blue denotes exact-target removal, and orange denotes the 70% sequence-deduplicated dataset.

**Task 2: molecule-conditioned synthesizability projection**

The Task 2 dataset was constructed as a molecule-only corpus without protein information. Molecules from the curated Task 1 dataset were pooled with eligible compounds from ChEMBL 36. ChEMBL records were required to have activity values below 10 μM, assay confidence scores of at least 8 and mappings to reviewed UniProt proteins. Molecular structures were converted to canonical SMILES, and only single-component molecules with SMILES strings no longer than 100 characters were retained. Salts and mixtures containing disconnected components (.) were excluded, and the allowed elements were restricted to C, H, N, O, S, P, F, Cl, Br and I.

Structures were deduplicated at the canonical-SMILES level. When several records described the same molecule, quantitative activity measurements were prioritized over categorical annotations. Among quantitative records, the lowest nanomolar activity value was retained. Because part of the Task 2 molecular pool was inherited from Task 1, corresponding datasets were prepared for both the exact-target-removal and 70% MMseqs2 sequence-deduplication settings. Each setting contained approximately $5.6 \times 10^5$ eligible molecular records and $2.1 \times 10^5$ unique molecules (**Supplementary Fig. 2**).

Sequence-based target deduplication had little effect on the chemical composition of the resulting molecule-only corpus. In both settings, the mean molecular weight was approximately 435 Da, the mean QED was 0.51, the mean heavy-atom count was 30.8, the mean cLogP was 3.84 and the mean topological polar surface area was approximately 87 $Å^2$. The near-complete overlap of these distributions indicates that the stricter protein-sequence separation did not introduce a substantial physicochemical shift into Task 2. The resulting molecules served as inputs for synthesizability projection, in which OmniSyn generated structurally related products together with explicit reaction trajectories.

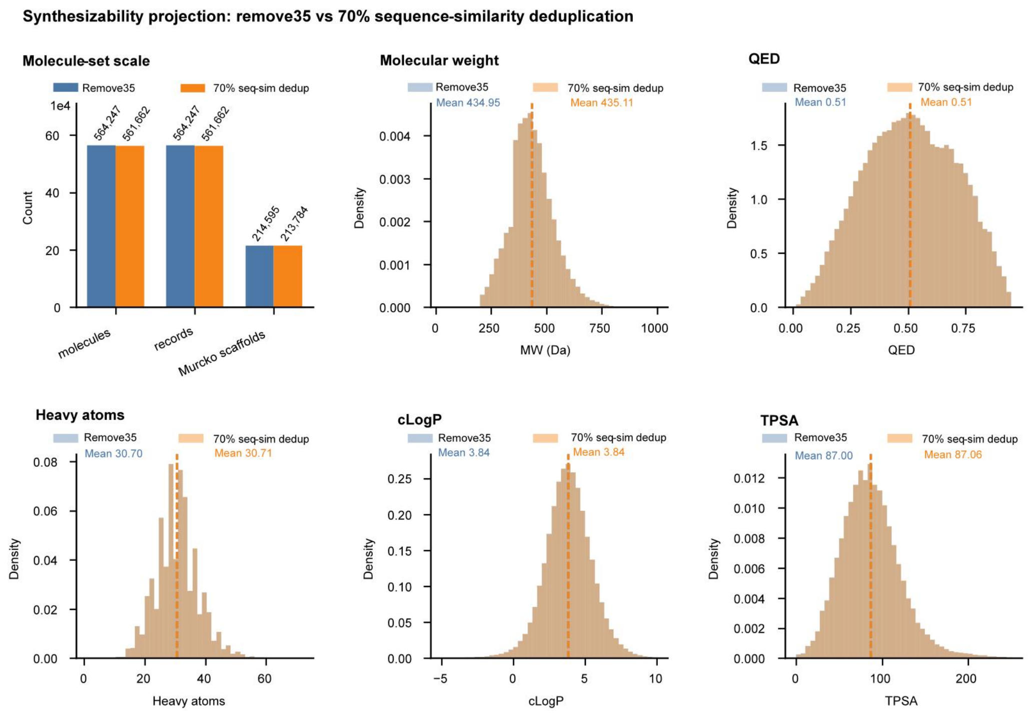


**Supplementary Fig. 2 | Composition and physicochemical properties of the Task 2 dataset. a**, Numbers of input molecular records, retained records and unique molecules in the exact-target-removal dataset (blue) and the dataset associated with 70% protein-sequence deduplication using MMseqs2 (orange). **b-f**, Distributions of molecular weight (**b**), quantitative estimate of drug-likeness (QED; **c**), heavy-atom count (**d**), calculated octanol-water partition coefficient (cLogP; **e**) and topological polar surface area (TPSA; **f**). Distributions are shown for the exact-target-removal and 70% sequence-deduplicated settings. Dashed vertical lines indicate the corresponding means.

**Task3: Hit to lead(H2L) optimization**

The Task 3 dataset was constructed from experimentally annotated chemical series in ChEMBL 36 and target-associated compounds in Papyrus. ChEMBL records provided explicit series or assay groupings, whereas comparable series annotations were unavailable for Papyrus. We therefore preserved the ChEMBL groupings and inferred putative Papyrus series by target-wise chemical clustering. For each series, a maximum common substructure (MCS) was extracted as the molecular context. Each training example comprised a protein sequence, the corresponding series-level MCS and a complete member molecule as the prediction target.

For ChEMBL 36, molecules were grouped by their reported series or assay identifiers. We retained exact $K_i$, $K_d$, $IC_{50}$ and $EC_{50}$ measurements reported in nanomolar units, with values between 0 and 100,000 nM, assay confidence scores of at least 8 and mappings to reviewed UniProt proteins. Structures were restricted to single-component canonical SMILES no longer than 100 characters and containing only C, H, N, O, S, P, F, Cl, Br or I. Within each series, the most frequently reported activity type was retained, and duplicate canonical SMILES were resolved by selecting the strongest available measurement.

For Papyrus, molecules were first grouped by protein target and deduplicated at the canonical-SMILES level. Because Papyrus lacks consistent series annotations, putative series were inferred using 2,048-bit ECFP4 fingerprints. Molecules were clustered with the Butina algorithm using a distance cutoff of 0.4, corresponding to a Tanimoto similarity threshold of 0.6. For targets containing more than 3,000 molecules, sphere-exclusion clustering with the same cutoff was used to reduce computational cost. Each resulting cluster was assigned a target-specific pseudo-series identifier.

MCS extraction was performed using RDKit with atom-identity and bond-order matching, ring-to-ring matching and complete-ring preservation. The MCS threshold was decreased from 1.0 to 0.4 in increments of 0.1, and the highest threshold producing an acceptable scaffold was retained. A valid MCS was required to occur in at least 80% of the corresponding series and to cover at least 33% of the atoms in each retained molecule. ChEMBL series containing more than 80 molecules and Papyrus clusters containing more than 50 molecules were first subdivided by Bemis-Murcko scaffold. Singleton Papyrus clusters were represented by their Bemis-Murcko scaffold, whereas two-member clusters were processed by direct MCS extraction. Acyclic singletons without a valid scaffold were excluded.

ChEMBL and Papyrus series were subsequently merged, with ChEMBL series identifiers and MCS assignments preserved. Papyrus-derived records overlapping ChEMBL at the level of UniProt identifier and canonical SMILES were removed.

The primary Task 3 dataset contained 562,036 series-member records spanning 4,847 proteins, 329,402 chemical series, 114,835 MCS scaffolds and 392,546 unique molecules. After MMseqs2 filtering at 70% protein-sequence similarity, 547,459 members, 4,755 proteins, 320,247 series, 113,015 MCS scaffolds and 385,149 molecules remained. Binder annotations constituted the largest activity class, decreasing from 217,503 to 213,643 records after filtering, followed by $IC_{50}$ measurements (150,843 to 147,089), $pIC_{50}$ measurements (134,448 to 128,438) and $K_i$ measurements (53,089 to 52,162). The mean molecular weight changed only marginally, from 425.24 to 425.06 Da, and the mean QED remained 0.52. Thus, sequence-level deduplication reduced potential protein overlap while preserving the scale and chemical composition of the Task 3 corpus.

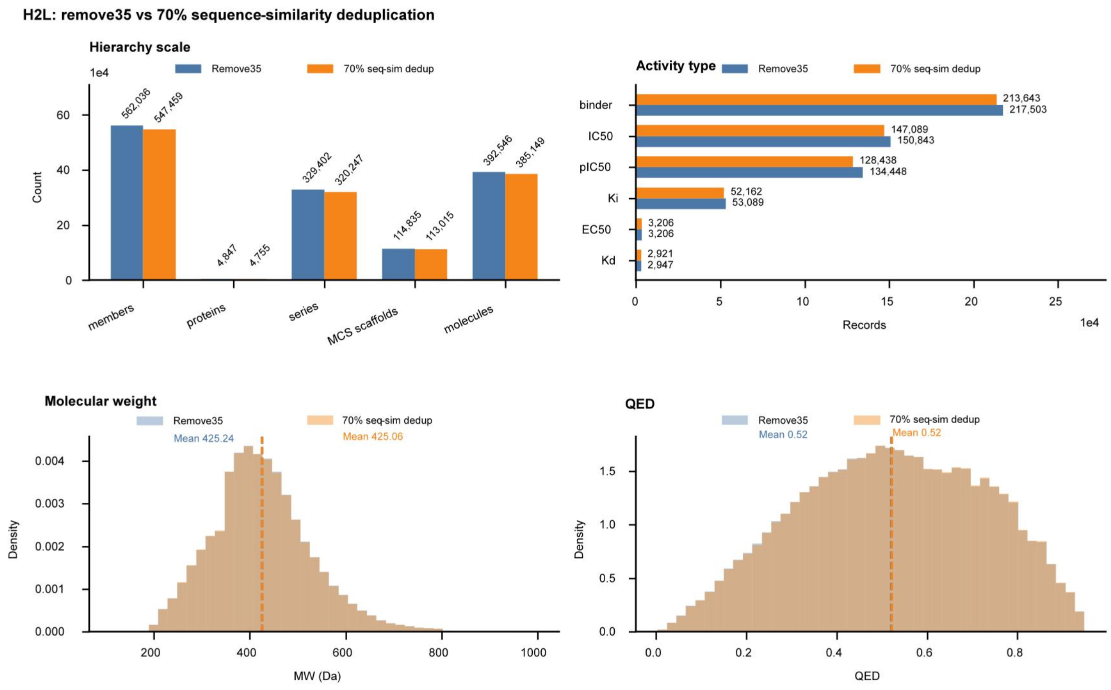

**Supplementary Fig.** 3 | Composition of the Task 3 dataset after target- and sequence-level deduplication. **a,** Hierarchical composition of the Task 3 H2L dataset under exact removal of the 35 MolGenBench targets and the more stringent MMseqs2-based 70% sequence-similarity filter. Bars show the numbers of series-member records, protein targets, chemical series, distinct MCS variants and protein–series contexts. **b,** Numbers of records associated with each retained activity type, including binder, Ki, Kd, IC50 and EC50 or pIC50. **c,d,** Density distributions of molecular weight (**c**) and QED (**d**) under the two deduplication settings. Dashed vertical lines indicate the corresponding means. Blue denotes exact removal of the 35 MolGenBench targets, and orange denotes MMseqs2-based sequence deduplication at a 70% similarity threshold.

### Supplementary Information of physicochemical distributions generated by different models

We compared the physicochemical distributions of all de novo molecules generated for the 35 unseen MolGenBench targets with those of the corresponding reference active ligands (**Supplementary Figs. 4,5**). Across $Fsp^3$, hydrogen-bond acceptor count, molecular weight, aromatic-ring count, rotatable-bond count and logP, OmniSyn consistently reproduced both the central tendencies and the broader distributional profiles of the reference set. This agreement indicates that OmniSyn generated molecules within a physicochemical regime characteristic of known active ligands rather than concentrating on a restricted region of chemical space.

Several baseline models matched the reference distribution for individual properties but showed pronounced shifts in others, including enrichment for lower-molecular-weight compounds or altered aromaticity, flexibility and lipophilicity. By contrast, OmniSyn maintained comparatively consistent agreement across all six properties. These results complement the aggregate molecular-quality metrics by showing that OmniSyn preserves reference-like physicochemical diversity across the complete set of unseen MolGenBench targets.

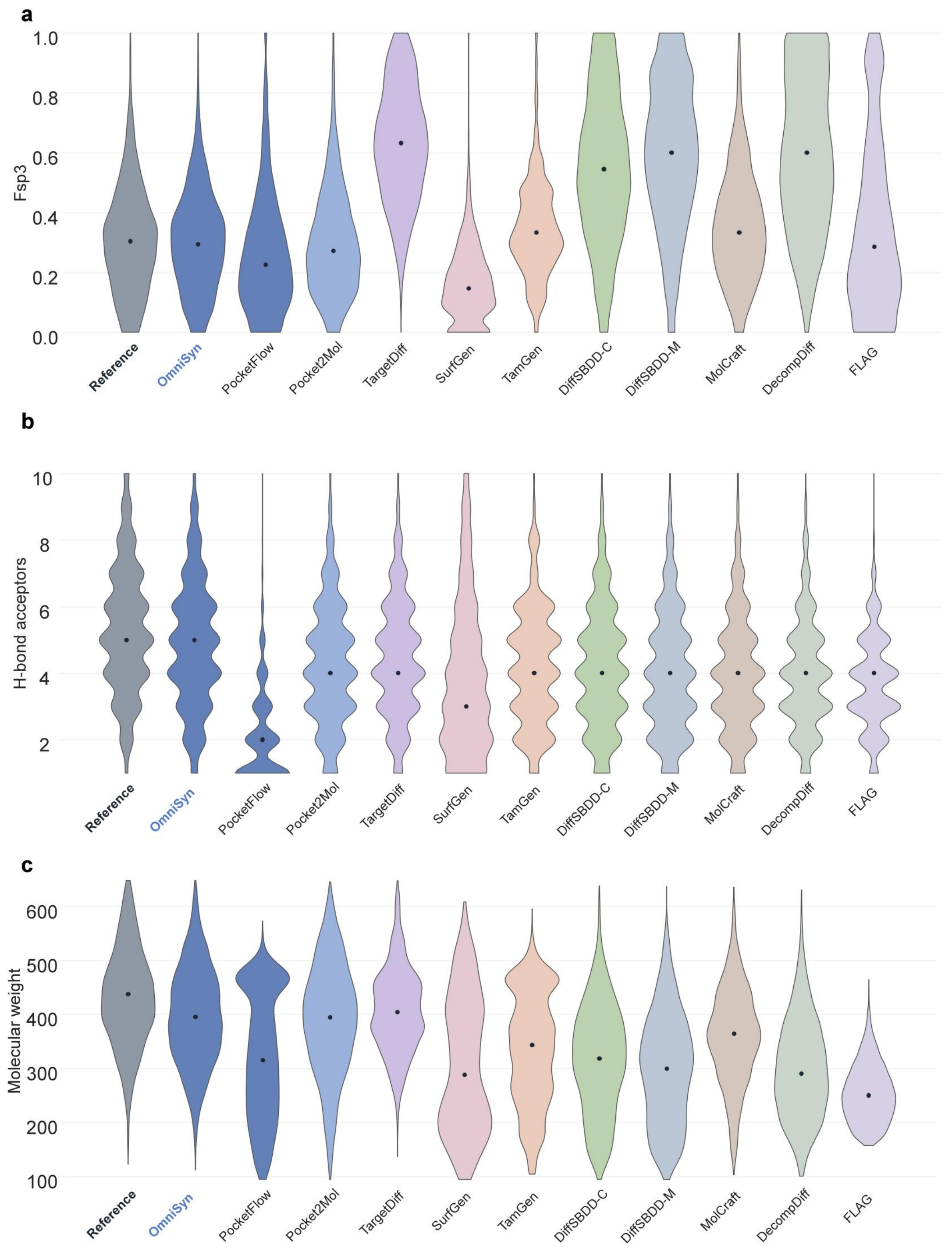


**Supplementary Fig. 4** | OmniSyn reproduces the physicochemical distributions of reference active ligands in de novo generation. Violin plots compare the distributions of physicochemical properties for reference active ligands and molecules generated by OmniSyn and the baseline models. **a,** Fraction of $sp^3$-hybridized carbon atoms ($Fsp^3$). **b,** Number of hydrogen-bond acceptors. **c,** Molecular weight.

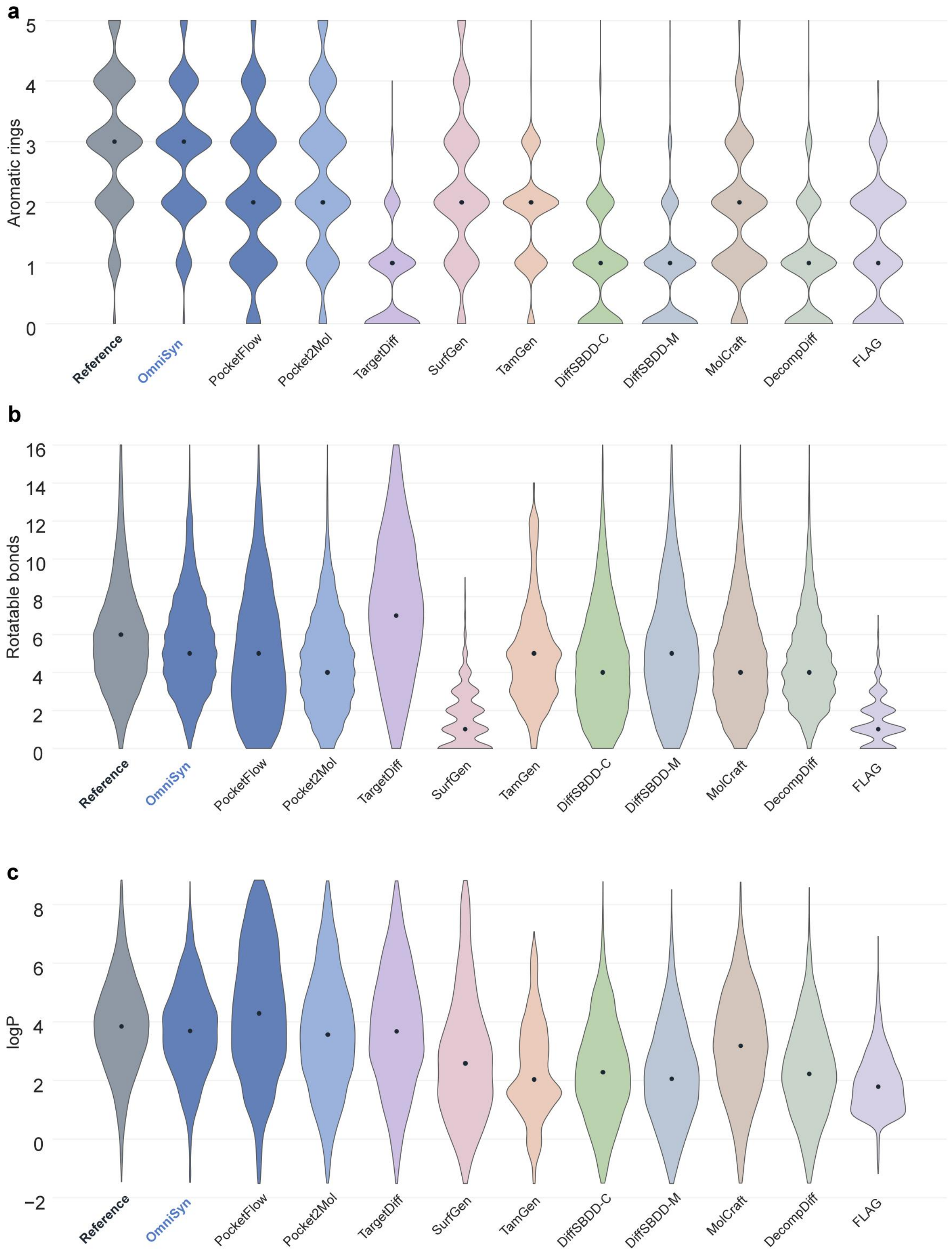


**Supplementary Fig. 5** | OmniSyn reproduces the physicochemical distributions of reference active ligands in de novo generation. Violin plots compare the distributions of physicochemical properties for reference active ligands and molecules generated by OmniSyn and the baseline models. **a,** Number of aromatic rings. **b,** Number of rotatable bonds. **c,** Octanol-water partition coefficient (logP). Violin widths represent kernel-density estimates, and black points denote the median of each distribution. Molecules were evaluated using the standardized MolGenBench protocol.

**Supplementary Information for MolGenBench evaluation results after deduplicating by similarity**

To assess whether OmniSyn's performance could be attributed to close sequence overlap between benchmark targets and the training data, we constructed a sequence-deduplicated MolGenBench subset using MMseqs2[4]. Applying a stricter deduplication procedure than previous models, we queried the 35 'unseen' MolGenBench protein targets against the OmniSyn training data. Any training proteins exhibiting 70% or higher sequence identity to these benchmark targets were entirely removed from our training set, alongside their associated ligand data." This procedure retained 35 sequence-deduplicated protein targets for evaluation. All methods were evaluated on this identical 35 'unseen' targets using the original MolGenBench protocol, including the generation budget, molecular standardization, chemical filters, reference active molecules and definitions of hit fraction, hit rate and Target-Aware score. Metrics were calculated for each target and aggregated across three sampling rounds.

OmniSyn retained strong molecular quality and synthetic accessibility under this more stringent evaluation (**Supplementary Figs. 6a,b**). It achieved a validity of 0.951 and the highest normalized SA score (0.795), atom-type score (0.947), ring-type score (0.922) and functional-group score (0.740), while maintaining competitive QED and diversity. OmniSyn also obtained the highest scaffold-level chemical-filter pass fraction (0.443) and the second-highest SMILES-level pass fraction (0.588). These results indicate that the molecular quality observed in the full benchmark was preserved after excluding targets closely related to those represented during training.

Active-molecule recovery remained similarly robust (**Supplementary Fig. 6c**). OmniSyn achieved the highest scaffold- and SMILES-level hit fractions, at 2.027% and 0.197%, respectively, together with the highest SMILES hit rate of 0.421%. Its scaffold hit rate remained strong at 5.516%, although PocketFlow attained a higher value for this individual metric. Target-aware analysis further distinguished OmniSyn from the baseline models (**Supplementary Fig. 7**). At the SMILES level, only 28% of targets occupied the lowest TA-score interval of 0-1, compared with 92-100% for most baselines; 60% and 10% of targets instead reached the 10-100 and greater-than-100 intervals, respectively. At the scaffold level, 64% of targets fell within the TA-score interval of 1-10. Collectively, these results show that OmniSyn preserves molecular quality, active-compound recovery and target specificity on sequence-deduplicated targets, supporting generalization beyond close protein homologues represented in the training data.

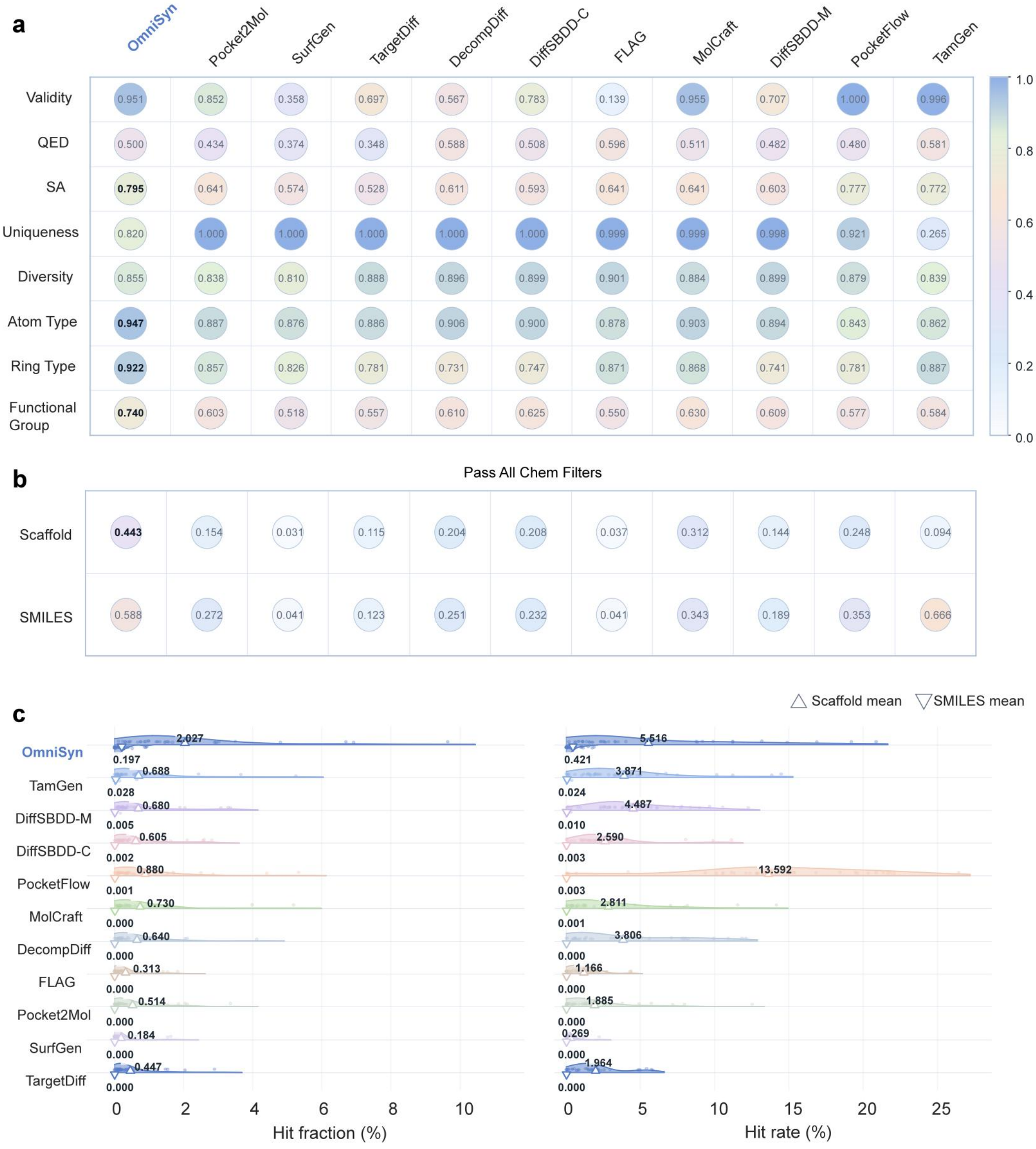

**Supplementary Fig. 6** | Molecular quality and active-molecule recovery on the sequence-deduplicated MolGenBench benchmark. Results are shown for the MolGenBench evaluation subset defined using a 70% protein-sequence similarity threshold against the OmniSyn training data. **a,** Comparison of validity, quantitative estimate of drug-likeness (QED), synthetic accessibility (SA), uniqueness, molecular diversity, and the agreement of atom-type, ring-type and functional-group distributions with the reference molecules. Circle colour and the values shown indicate the corresponding metric scores. **b,** Fractions of generated scaffolds and molecules, represented as SMILES, that pass all MolGenBench chemical filters. **c,** Target-wise distributions of scaffold- and SMILES-level hit fractions (left) and hit rates (right). Upward- and downward-pointing triangles indicate the across-target means for scaffolds and SMILES, respectively.

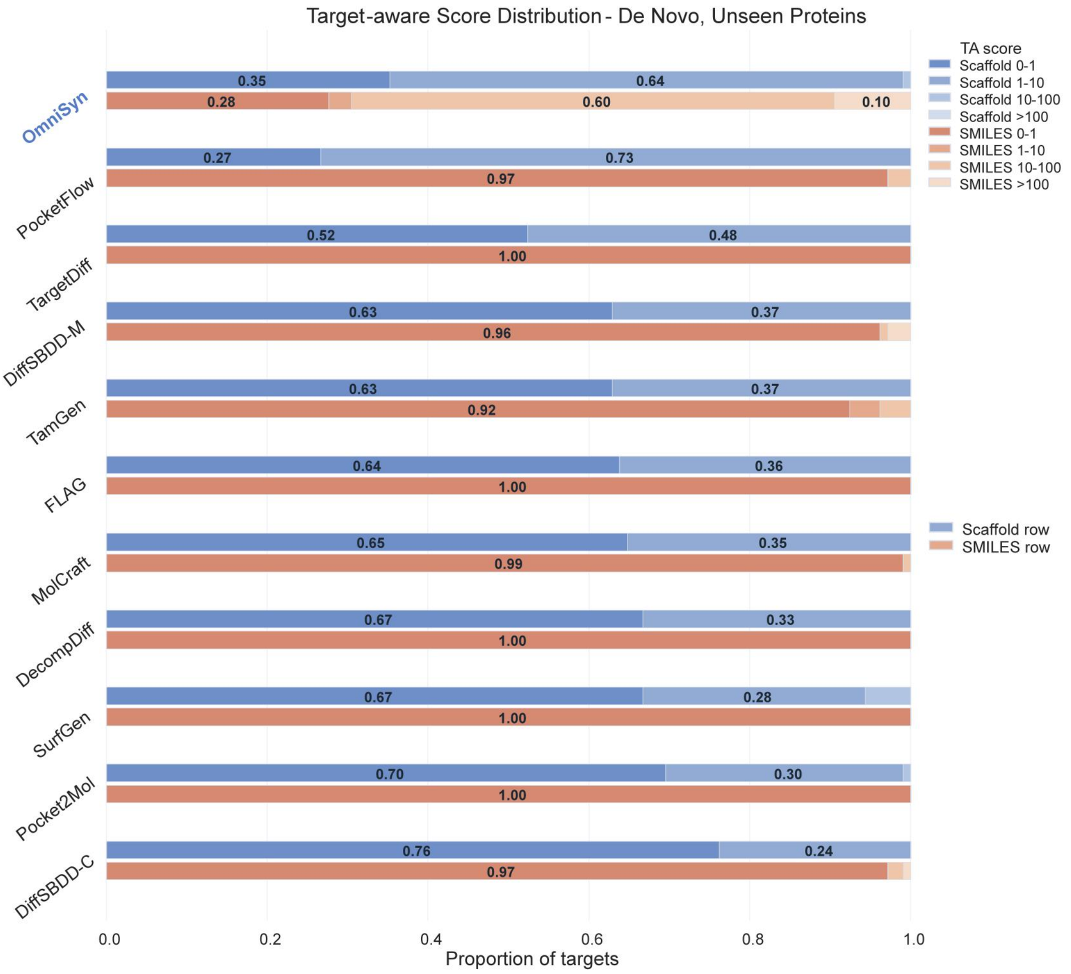


**Supplementary Fig. 7** | Target-aware recovery on the sequence-deduplicated MolGenBench benchmark. Distribution of scaffold and SMILES-level Target-Aware (TA) scores across the MolGenBench evaluation targets retained using a 70% protein-sequence similarity threshold against the OmniSyn training data. For each method, the blue and orange stacked bars represent scaffold- and SMILES-level results, respectively. Bar segments indicate the proportions of targets with TA scores of 0-1, 1-10, 10-100 and greater than 100. Values within the segments denote the corresponding proportions of targets.

### Supplementary Information for detailed PoseBusters evaluation of generated molecules

We performed a detailed PoseBusters[5] analysis to identify the structural factors underlying the aggregate three-dimensional validity reported in the main text. Following the MolGenBench protocol, generated molecules were embedded using RDKit, redocked to their corresponding protein targets with AutoDock Vina and evaluated using individual PoseBusters checks. These checks assess molecular loading and sanitization, InChI convertibility, molecular connectivity, bond lengths and angles, intramolecular clashes, aromatic-ring and double-bond planarity, internal energy, and protein-ligand distance and volume-overlap constraints. The aggregate pass-all criterion required a molecule to satisfy every individual check simultaneously.

In the de-novo setting, OmniSyn showed consistently high pass fractions across all categories **(Supplementary Fig. 8a**). Approximately 95% of generated molecules passed the molecular-validity and connectivity checks, as well as the bond-length and bond-angle criteria. High pass fractions were also observed for intramolecular clashes (0.93), aromatic and double-bond planarity (approximately 0.95), and internal energy (0.91). Protein-ligand spatial checks remained similarly favorable, with pass fractions of 0.91–0.94 for the distance and overlap criteria. Consequently, OmniSyn achieved the highest aggregate pass-all fraction of 0.87, indicating that its three-dimensional validity was not driven by performance on only a subset of structural checks.

Performance was further improved in the H2L setting (**Supplementary Fig. 8b**). OmniSyn-H2L achieved pass fractions approaching 1.00 for molecular validity, connectivity, bond geometry and planarity, together with pass fractions of 0.99 for intramolecular clashes and 0.96 for internal energy. The protein–ligand spatial compatibility checks reached 0.98-0.99, resulting in an aggregate pass-all fraction of 0.95, substantially exceeding those of the evaluated H2L baselines. Thus, conditioning generation on an existing molecular context did not introduce detectable geometric distortions or steric liabilities into the optimized molecules.

Together, these results show that OmniSyn generates molecules that remain chemically valid, internally well-formed and spatially compatible with the target environment after independent conformer generation and redocking. The consistently high performance across individual PoseBusters criteria further supports the three-dimensional plausibility of both de novo and H2L outputs, despite the absence of explicit protein-pocket structures during generation.

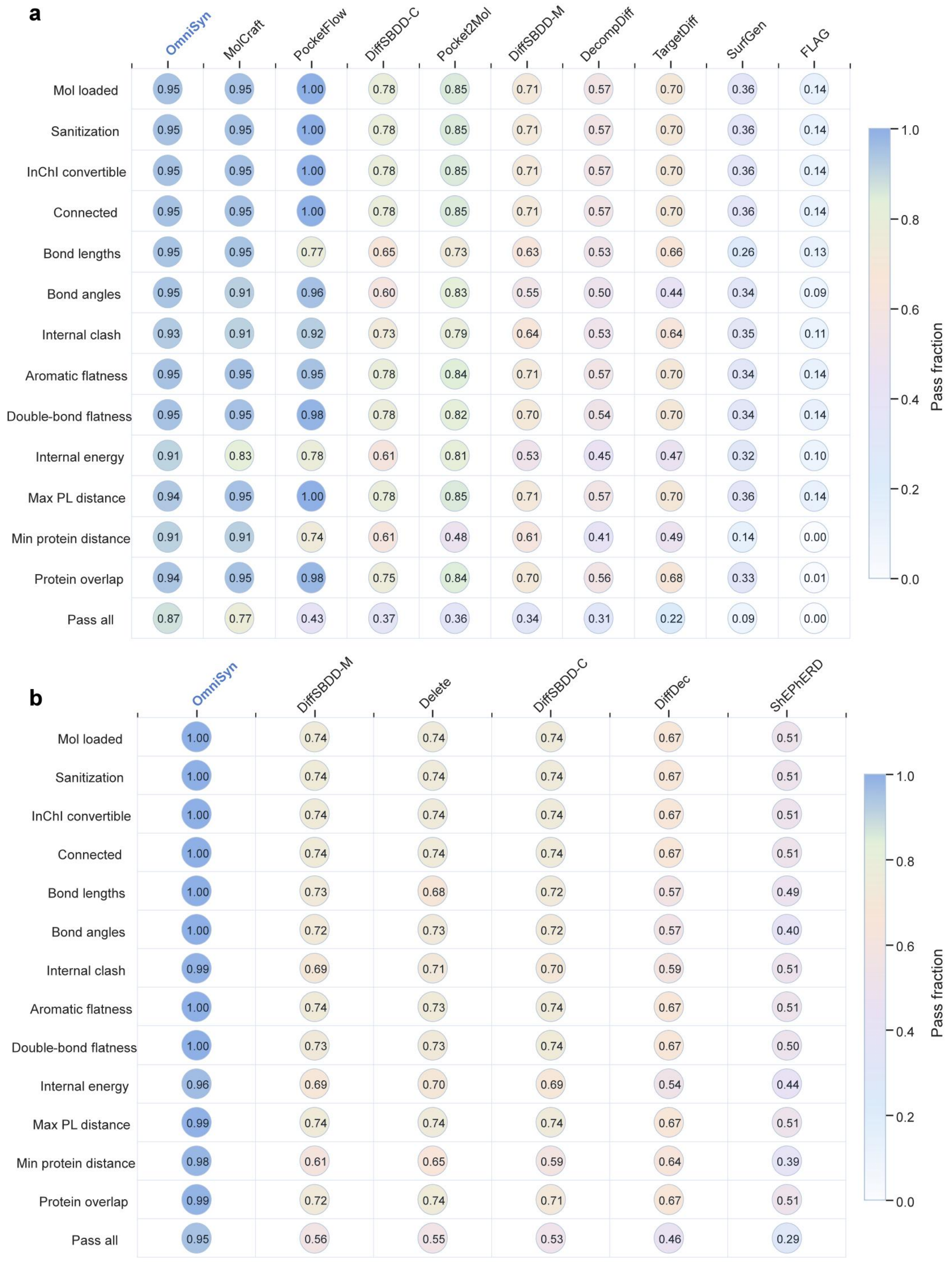

**Supplementary Fig. 8** | Detailed PoseBusters evaluation of generated molecules. Pass fractions for individual PoseBusters checks and the aggregate pass-all criterion in **a,** de novo generation and **b,** H2L optimization. The evaluation covers molecular validity and connectivity, bond geometry, intramolecular clashes and planarity, internal energy, and protein–ligand spatial compatibility. Values and circle colours denote the fraction of molecules passing each check, with higher values indicating better performance. Molecules were evaluated after RDKit conformer generation and AutoDock Vina redocking using the MolGenBench protocol.

### Supplementary Information for Additional example of synthesizability projection by OmniSyn

We examined an additional example to assess whether OmniSyn could improve retrosynthetic feasibility while preserving the structural and predicted binding characteristics of the input molecule (**Supplementary Fig. 9**). The input compound adopted a plausible pose within the target pocket and achieved a Vina docking score of -7.135 kcal/mol (**Supplementary Fig. 9a**). However, AiZynthFinder failed to identify a viable retrosynthetic solution, illustrating that favourable predicted binding does not necessarily imply synthetic tractability.

OmniSyn converted this compound into a closely related analogue with an ECFP4 similarity of 0.800 (**Supplementary Fig. 9b**). The projected molecule occupied a comparable region of the binding pocket and retained the principal interaction pattern of the input compound. Its Vina docking score improved to −8.126 kcal/mol, and AiZynthFinder successfully identified a retrosynthetic solution. Thus, improved retrosynthetic feasibility was achieved without substantial departure from the input chemical structure or loss of predicted binding compatibility.

The projected molecule was also accompanied by a model-derived, template-level synthesis route assembled from purchasable building blocks (**Supplementary Fig. 9c**). Reaction 68 formally combines a methoxybenzenesulfonyl chloride building block (94173), a branched amino-acid building block (24078) and a pyridine-carbaldehyde building block (64275) through an annotated C-N bond-forming transformation. This route represents a reaction-template-level synthesis hypothesis rather than an experimentally optimized protocol. Together, this example provides an independent illustration of OmniSyn converting a retrosynthetically unresolved molecule into a structurally similar analogue with improved docking performance, external retrosynthetic solvability and an explicit model-derived reaction trace.

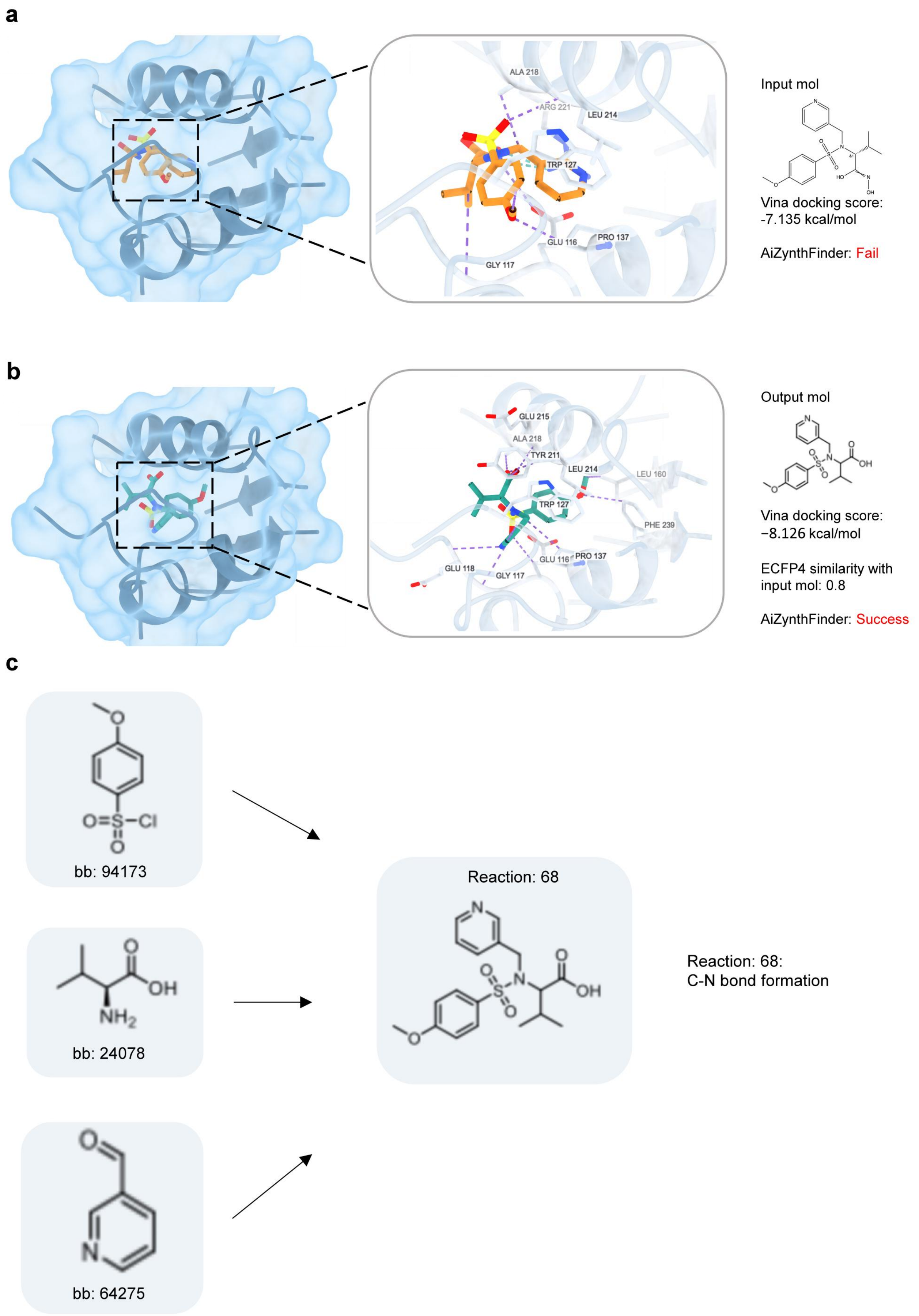
a
ALA 218
ARG 221
LEU 214
TRP 127
GLU 116
PRO 137
GLY 117
Input mol
Vina docking score:
-7.135 kcal/mol
AiZynthFinder: Fail
b
GLU 215
ALA 218
TYR 211
LEU 214
LEU 160
TRP 127
PHE 239
GLU 116
PRO 137
GLU 118
GLY 117
Output mol
Vina docking score:
−8.126 kcal/mol
ECFP4 similarity with
input mol: 0.8
AiZynthFinder: Success
c
bb: 94173
bb: 24078
bb: 64275
Reaction: 68
Reaction: 68:
C-N bond formation

**Supplementary Fig. 9** | Additional example of synthesizability projection by OmniSyn. **a,** Predicted binding pose of the input molecule and an enlarged view of its interactions within the protein-binding pocket. The input molecule achieved a Vina docking score of -7.135 kcal/mol, but AiZynthFinder failed to identify a synthetic route. **b,** Predicted binding pose and protein-ligand interactions of the projected molecule. The projected molecule retained an ECFP4 similarity of 0.800 to the input, improved the Vina docking score to -8.126 kcal/mol and was successfully solved by AiZynthFinder. **c,** Template-level synthetic route proposed for the projected molecule from purchasable building blocks. The route combines building blocks 94173, 24078 and 64275 through Reaction 68, annotated as C-N bond formation. Dashed purple lines in a and b indicate predicted protein-ligand interactions.

### Supplementary Information for Task-dependent expert routing with PLIM

We examined whether PLIM allocated distinct computational pathways to the three generation tasks (**Supplementary Fig. 10**). De novo generation most frequently selected E0 (0.29), with additional routing to E5 and E7, whereas synthesizability projection preferentially selected E6 (0.22) and distributed substantial routing across E2-E4. H2L optimization also favoured E0 (0.21), but showed a more distributed profile with increased selection of E7. These partially shared yet distinct routing patterns indicate that PLIM adapts expert utilization to task-specific inputs while retaining computational components across related design settings.

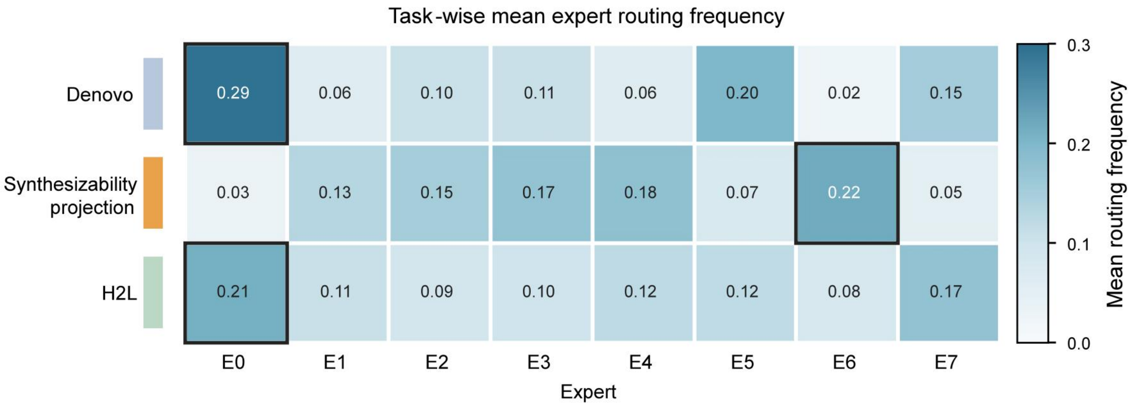


**Supplementary Fig. 10** | Task-dependent expert routing within PLIM. Heat map showing the mean top-1 routing frequency of experts E0-E7 for de novo generation, synthesizability projection and H2L optimization. Routing frequencies were averaged across valid pair positions, input samples and the four PairFormer-MoE blocks. Cell values and colour intensity indicate how frequently each expert was selected, and black outlines identify the most frequently selected expert for each task.

**Supplementary Information for SAE analysis**

To examine how chemical and synthesis information is distributed across the final OmniSyn checkpoint, we fitted TopK SAEs ($K$ = 32) to the PLIM conditioning representation and to reaction-decoder layers 0, 5 and 11. We selected 500 SAE features per representation for enrichment testing. For each selected feature, the preferred task was defined by the highest activation frequency in the full scan. Up to 30 positive top-activating states were then selected within that task, and the background was restricted to the same task. Chemical enrichment used unique final products defined by isomeric canonical SMILES. For decoder representations, token-type, building-block and reaction-template enrichment was evaluated at decoder-state level. One-sided hypergeometric $P$ values were adjusted across all tested feature-attribute associations within each representation using the Benjamini-Hochberg procedure.

No conditioning-representation feature showed a significant chemical or route association after multiple-testing correction. In the decoder, 399, 979 and 1,128 feature-attribute associations remained significant at layers 0, 5 and 11, respectively. Among the 500 features tested at each layer, 48, 176 and 253 features had at least one significant association with a molecular substructure or Bemis-Murcko scaffold, whereas 272, 457 and 450 features had at least one significant association with token type, reaction-template identity or building-block identity. Chemical and route counts are not mutually exclusive because one feature can be associated with both classes of attribute. The layer-0 and layer-5 enrichment landscapes are shown in **Supplementary Fig. 11**, with the layer-11 landscape shown in **Fig. 6d**. Chemical associations increased across decoder depth, whereas route associations were abundant at all three layers and peaked at layer 5.

At layer 0, representative chemistry-associated features were enriched for carboxylic acid and benzimidazole-linked motifs (**Supplementary Fig. 12 a,b**). Other layer-0 features were associated with individual building-block identities or with reaction templates annotated as tert-butoxycarbonyl(Boc) deprotection and five-membered heterocycle formation (**Supplementary Fig. 12c-f**). These results show that associations with both the final product and recorded route decisions were already detectable in the earliest decoder layer.

The intermediate and final decoder layers contained further chemistry- and route-associated features. Layer-5 examples included a fused nitrogen-containing heterocycle, an ester functionality, distinct building-block identities and reaction templates annotated as urea or pyrazole formation (**Supplementary Fig. 13**). Layer-11 examples included pyrimidinedione and halogen-associated chemistry, distinct building-block identities, alkynyl C-C coupling and sulfonyl chloride activation (**Supplementary Fig. 14**). In the route panels, the displayed molecules are final products reached by high-activation states carrying the enriched decision; they are not depictions of the building block or reaction template itself. Together, the layer-resolved analyses indicate that sparse decoder features are associated with both final-product chemistry and synthesis-path decisions. These associations are descriptive and do not establish that intervening on an individual feature would causally control generation.

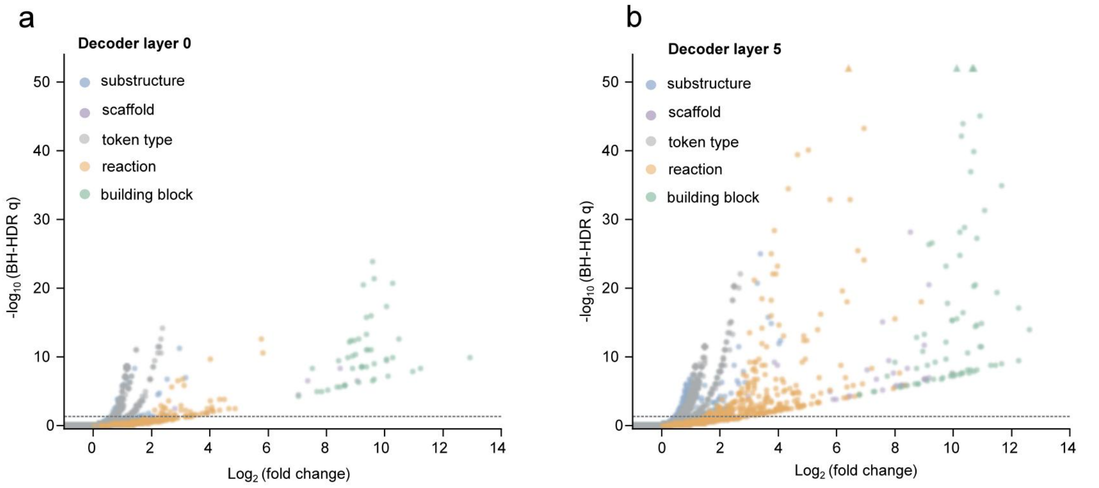


**Supplementary Fig. 11** | Layer-resolved enrichment landscapes for decoder SAE features. **a,b,** Enrichment results for decoder layers 0 and 5, respectively. Each point represents one tested association between an SAE feature and a molecular substructure, Bemis-Murcko scaffold, token type, reaction-template identity or building-block identity. The horizontal axis shows the base-2 logarithm of fold enrichment, and the vertical axis shows the negative base-10 logarithm of $q$, where $q$ is the Benjamini-Hochberg-adjusted one-sided hypergeometric $P$ value after correction across all tested feature-attribute associations within the indicated layer. The dashed line denotes $q = 0.05$; triangles denote associations above the displayed vertical range. Feature and background populations were matched by task. Chemical tests used unique isomeric canonical final products as the statistical unit, whereas route tests used decoder states; building-block and reaction-template tests were further restricted to states carrying a valid identity.

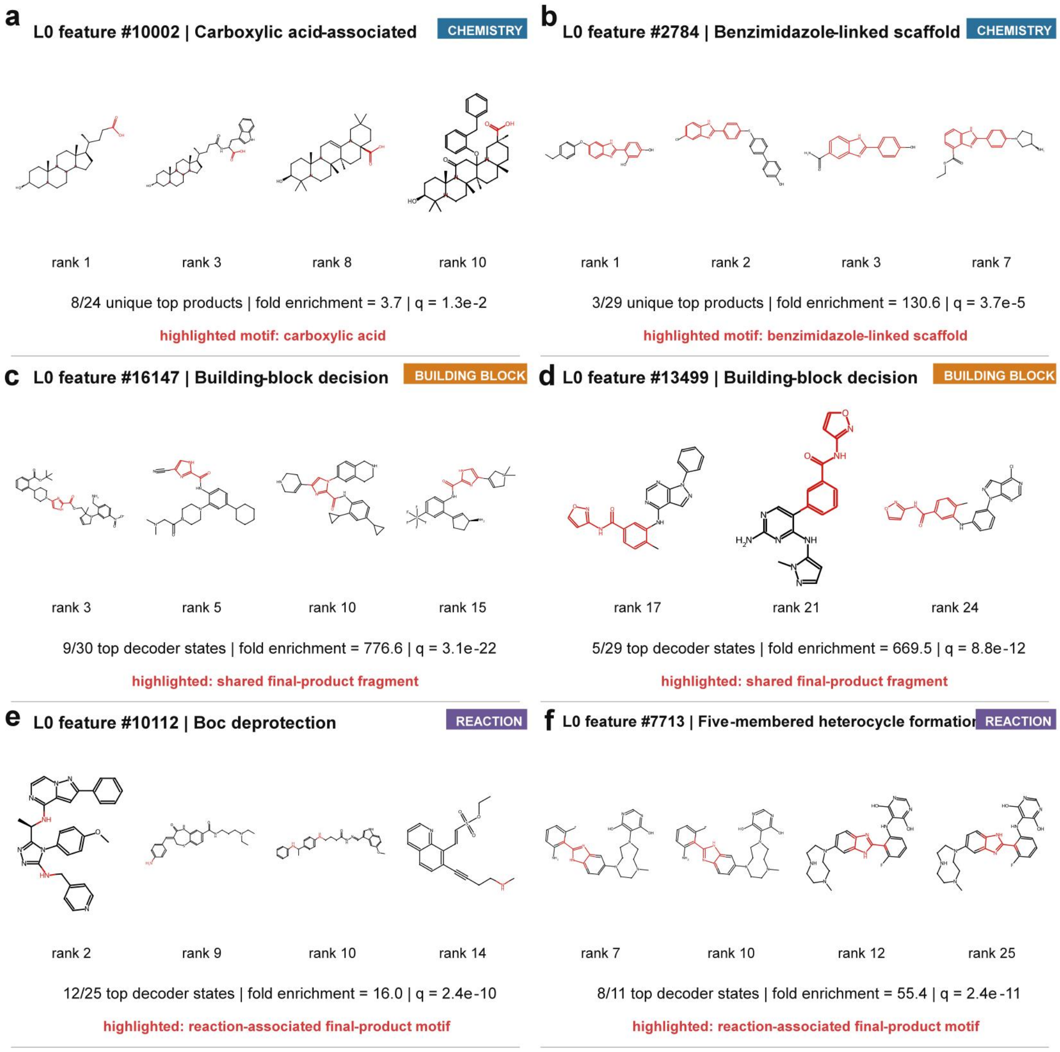


**Supplementary Fig. 12** | Representative layer-0 decoder SAE features associated with product chemistry and synthesis decisions. **a,b,** Features associated with final products enriched for carboxylic acid and a benzimidazole-linked scaffold, respectively. **c,d,** Features associated with distinct building-block identities. **e,f,** Features associated with reaction templates annotated as Boc deprotection and five-membered heterocycle formation, respectively. Molecules are distinct final products linked to task-matched highest-activation states; displayed ranks refer to the corresponding decoder-state ranks before final-product deduplication. For chemistry panels, fractions indicate motif-containing unique products among the feature-associated unique top products. For route panels, fractions indicate states carrying the specified decision among the feature-associated top decoder states. Fold enrichment and q values are from one-sided hypergeometric tests with Benjamini-Hochberg correction across all tested feature-attribute associations within the indicated layer. Highlighted atoms denote the enriched chemical motif, a shared final-product fragment among states carrying the enriched building-block decision, or a reaction-associated final-product motif.

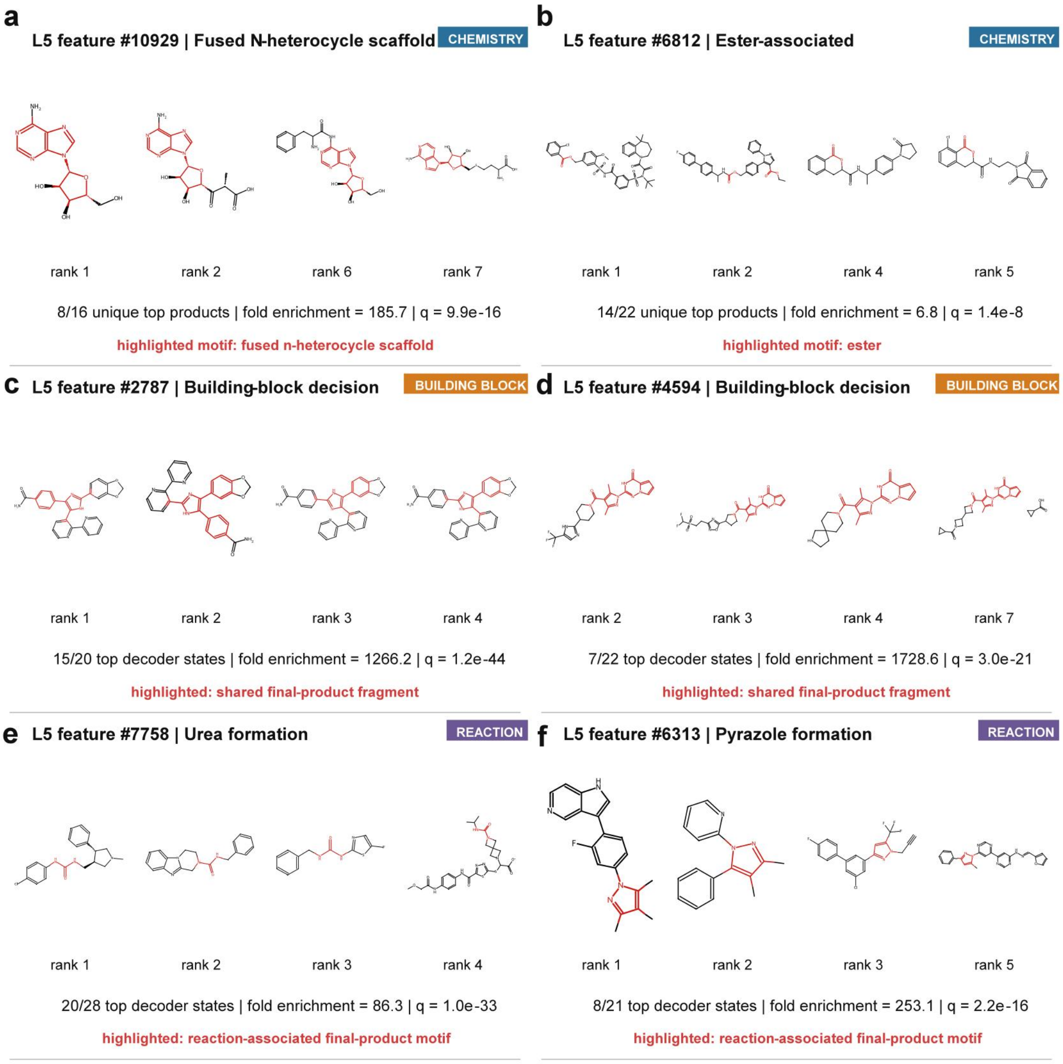


**Supplementary Fig. 13** | Representative layer-5 decoder SAE features associated with product chemistry and synthesis decisions. **a,b,** Features associated with final products enriched for a fused nitrogen-containing heterocycle and an ester functionality, respectively. **c,d,** Features associated with distinct building-block identities. **e,f,** Features associated with reaction templates annotated as urea and pyrazole formation, respectively.

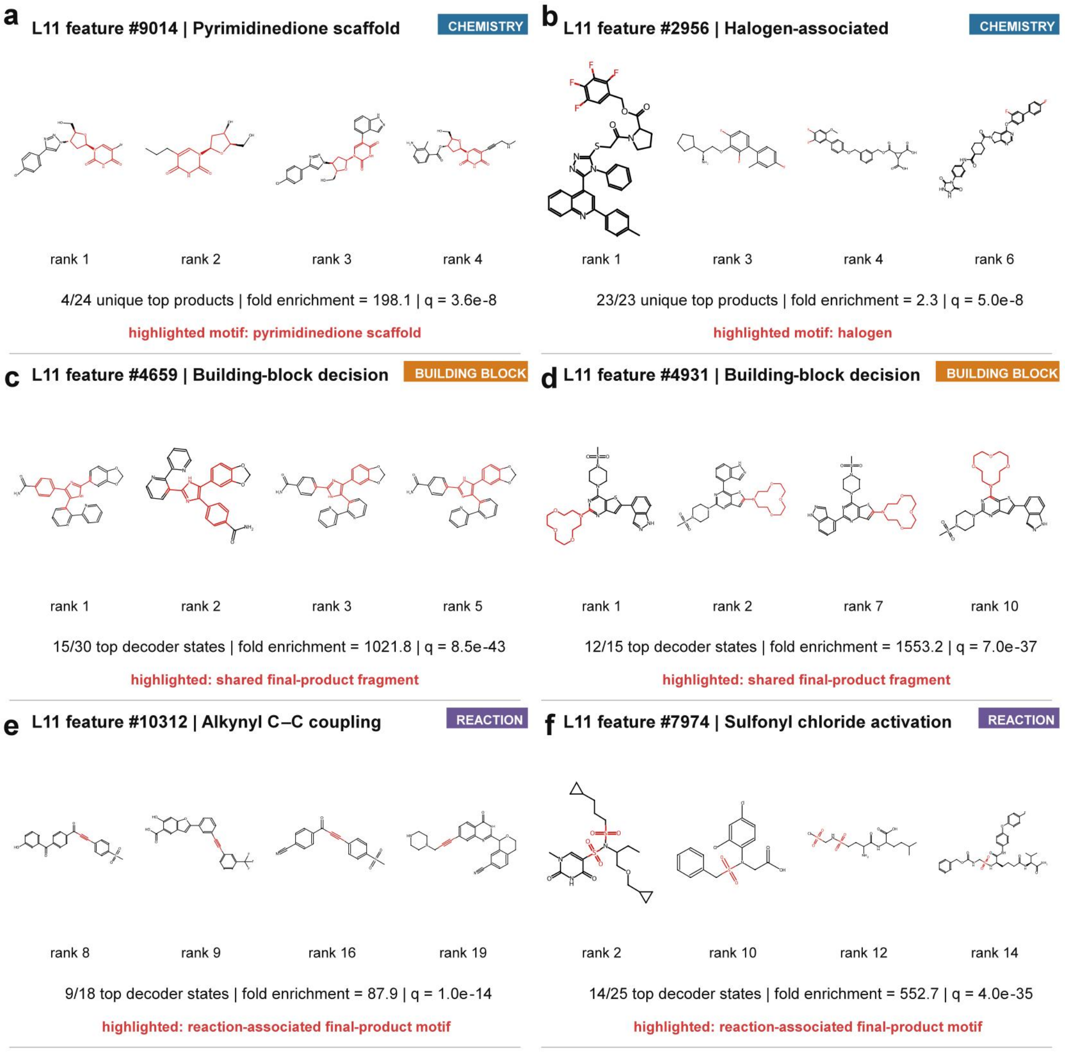


**Supplementary Fig. 14** | Representative layer-11 decoder SAE features associated with product chemistry and synthesis decisions. **a,b,** Features associated with final products enriched for a pyrimidinedione scaffold and halogen-containing motifs, respectively. **c,d,** Features associated with distinct building-block identities. **e,f,** Features associated with reaction templates annotated as alkynyl C-C coupling and sulfonyl chloride activation, respectively. Molecules are distinct final products linked to task-matched highest-activation states; displayed ranks refer to the corresponding decoder-state ranks before final-product deduplication. For chemistry panels, fractions indicate motif-containing unique products among the feature-associated unique top products. For route panels, fractions indicate states carrying the specified decision among the feature-associated top decoder states. Fold enrichment and q values are from one-sided hypergeometric tests with Benjamini-Hochberg correction across all tested feature-attribute associations within the indicated layer. Highlighted atoms denote the enriched chemical motif, a shared final-product fragment among states carrying the enriched building-block decision, or a reaction-associated final-product motif.

### Supplementary Information for case study in PD-L1

To examine whether OmniSyn generation was restricted to chemical neighbourhoods represented in the training data, we performed a case study on PD-L1. Known PD-L1 actives present in the OmniSyn training set were first removed, and the retained generated candidates were compared separately with the remaining held-out actives and PD-L1-associated training molecules. Many candidates were positioned above the diagonal in the similarity comparison (**Supplementary Fig. 15a**), indicating greater fingerprint similarity to a held-out active than to any target-associated training molecule. OmniSyn therefore sampled chemical space that extended beyond the nearest PD-L1 training examples while overlapping previously unseen active-like regions.

Two representative candidates illustrated this behavior (**Supplementary Fig. 15b, c**). Cases 1 and 2 showed higher maximum Tanimoto similarity to their corresponding held-out actives than to PD-L1-associated training molecules (0.66 vs 0.39 and 0.62 vs 0.36, respectively). Their predicted binding poses also showed spatial correspondence with the matched held-out actives. Although these computational comparisons do not establish experimental PD-L1 activity, they indicate that OmniSyn can generate candidates resembling unseen active chemistry without simply reproducing the closest target-associated training compounds.

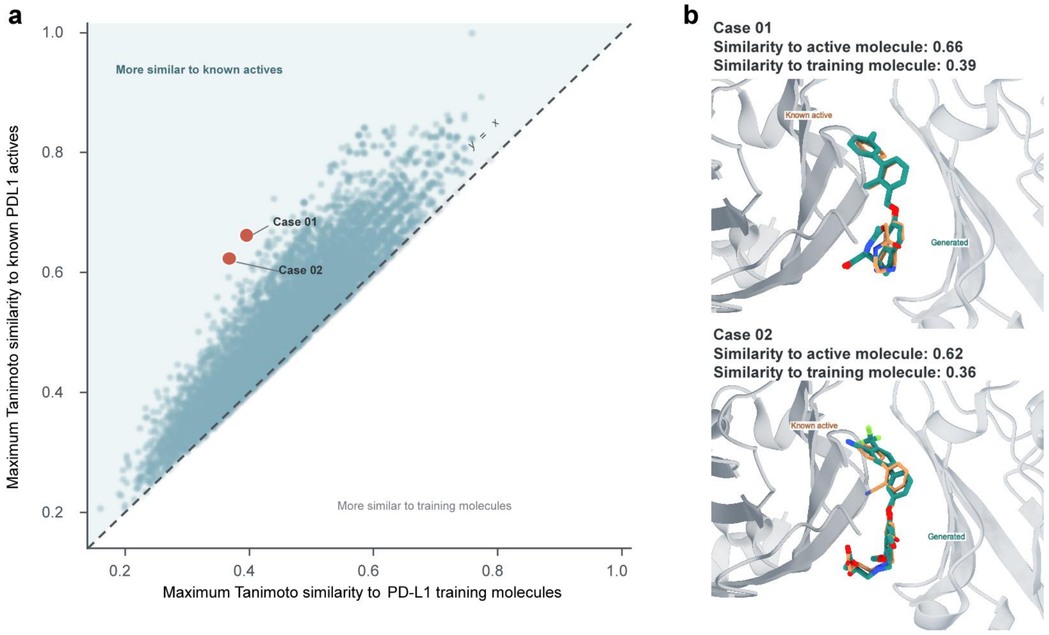

**Supplementary Fig. 15 | Recovery of held-out active-like chemical space for PD-L1**. **a,** Maximum molecular-fingerprint Tanimoto similarity of each retained OmniSyn-generated candidate to PD-L1-associated training molecules (x-axis) and held-out known PD-L1 actives (y-axis). The known-active reference set was constructed after removing all molecules present in the OmniSyn training data. Candidates were retained after scoring and filtering with PSICHIC-plus, Vina and DrugCLIP. Each point represents one generated molecule, and the dashed diagonal denotes equal similarity to the two reference sets. Points above the diagonal are more similar to a held-out known active than to any PD-L1-associated training molecule. Orange points indicate the representative candidates shown in b. **b,** Predicted binding poses of cases 1 and 2, with generated molecules shown in teal and the corresponding held-out actives shown in orange. Case 1 had maximum Tanimoto similarities of 0.66 and 0.39 to held-out actives and training molecules, respectively. The corresponding values for case 2 were 0.62 and 0.36.
